# Cohesin-mediated 3D contacts tune enhancer-promoter regulation

**DOI:** 10.1101/2024.07.12.603288

**Authors:** Philine Guckelberger, Benjamin R. Doughty, Glen Munson, Suhas S. P. Rao, Yingxuan Tan, Xiangmeng Shawn Cai, Charles P. Fulco, Joseph Nasser, Kristy S. Mualim, Drew T. Bergman, Judhajeet Ray, Evelyn Jagoda, Chad J. Munger, Andreas R. Gschwind, Maya U. Sheth, Anthony S. Tan, Saul Godinez Pulido, Namita Mitra, David Weisz, Muhammad Saad Shamim, Neva C. Durand, Ragini Mahajan, Ruqayya Khan, Lars M. Steinmetz, Masato T. Kanemaki, Eric S. Lander, Alexander Meissner, Erez Lieberman Aiden, Jesse M. Engreitz

## Abstract

Enhancers are key drivers of gene regulation thought to act via 3D physical interactions with the promoters of their target genes. However, genome-wide depletions of architectural proteins such as cohesin result in only limited changes in gene expression, despite a loss of contact domains and loops. Consequently, the role of cohesin and 3D contacts in enhancer function remains debated. Here, we developed CRISPRi of regulatory elements upon degron operation (CRUDO), a novel approach to measure how changes in contact frequency impact enhancer effects on target genes by perturbing enhancers with CRISPRi and measuring gene expression in the presence or absence of cohesin. We systematically perturbed all 1,039 candidate enhancers near five cohesin-dependent genes and identified 34 enhancer-gene regulatory interactions. Of 26 regulatory interactions with sufficient statistical power to evaluate cohesin dependence, 18 show cohesin-dependent effects. A decrease in enhancer-promoter contact frequency upon removal of cohesin is frequently accompanied by a decrease in the regulatory effect of the enhancer on gene expression, consistent with a contact-based model for enhancer function. However, changes in contact frequency and regulatory effects on gene expression vary as a function of distance, with distal enhancers (*e.g.*, >50Kb) experiencing much larger changes than proximal ones (*e.g.*, <50Kb). Because most enhancers are located close to their target genes, these observations can explain how only a small subset of genes — those with strong distal enhancers — are sensitive to cohesin. Together, our results illuminate how 3D contacts, influenced by both cohesin and genomic distance, tune enhancer effects on gene expression.

## Introduction

In mammalian genomes, spatiotemporal gene expression is largely driven by enhancers. However, how enhancers affect gene expression is not yet fully understood.

The current predominant model of how enhancers regulate genes involves physical contact in 3D — the classic “looping” model^1^ — which has been supported and refined over the last 30 years by numerous lines of evidence. Enhancers and promoters are often observed to form long-range 3D loops, the presence of which correlates with changes in gene expression across cell types^2–7^. Repositioning enhancers in the genome has been observed to alter both 3D chromatin interactions and gene expression^8–13^. Tethering factors that force the formation of chromatin loops can enhance gene expression^14–17^. Enhancer-gene regulatory interactions can be predicted by multiplying enhancer activity by enhancer-promoter 3D contact frequency (the Activity-by-Contact (ABC) model^18^). Together, these observations suggest that 3D contacts contribute to enhancer function.

Recent studies have revealed that key factors involved in the formation of such 3D loops and contacts include architectural proteins such as CTCF and cohesin, which orchestrate loop extrusion^19–22^. However, the depletion of these architectural proteins have minimal impacts on gene expression, despite the loss of loops and contact domains^23–26^. These observations, as well as other recent studies^27,28^, have raised questions regarding the looping model and the role of cohesin and CTCF in mediating these physical interactions. Alternative hypotheses have emerged. One proposed idea is that enhancer-promoter physical interactions may occur through other mechanisms, such as transcription factor affinity or compartmentalization^29,30^. Others suggest that while cohesin and CTCF are necessary for establishing initial contacts, they may not be essential for their maintenance^31^. Alternatively, it has been proposed that the requirement of cohesin/CTCF for mediating contacts may be specific to certain enhancer-promoter pairs, such as long-range or inducible physical interactions^12,17,32,33^.

One approach to resolve these questions would be to examine how altering 3D contacts, such as through depletion of CTCF or cohesin, changes the regulatory effects of individual enhancers on their target promoters. However, such experiments have been challenging because genes are usually regulated by multiple enhancers and each individual enhancer may only have a small effect on gene expression.

To address this challenge, we developed CRISPRi of regulatory elements upon degron operation (CRUDO) — a high-throughput and sensitive approach to measure enhancer effects on gene expression before and after cohesin depletion in an endogenous genomic context. We applied CRUDO to systematically perturb all candidate enhancers around 5 genes whose expression depend on cohesin, and compared the effects of enhancers on gene expression with their 3D contact frequencies in the presence and absence of cohesin. We find that changing 3D contact frequency quantitatively tunes the effects of enhancers on gene expression, and that distal enhancers (*e.g.*, >50Kb) are far more dependent on cohesin to facilitate both their 3D contacts and regulatory effects. Moreover, by exploring maps of enhancer-promoter regulatory interactions from the ABC model and from previous perturbation experiments, we find that most genes are not regulated by strong distal enhancers and consequently would not be expected to require cohesin for their expression. These data demonstrate that enhancer-promoter contact frequency quantitatively tunes enhancer-promoter regulation, that cohesin depletion selectively alters the regulatory effects of distal enhancers, and that such distal enhancers are rare in human gene regulation. Together, these observations could explain why cohesin depletion immediately alters the expression of only a small number of genes, and illuminate how the 3D architecture of the genome tunes gene expression.

### Predicting effects of cohesin depletion with the Activity-by-Contact model

To study how cohesin-mediated 3D contacts affect enhancer regulation, we first constructed a regulatory map of enhancers in a human colorectal cancer HCT-116 cell line engineered to express RAD21 (a major subunit of the cohesin complex) fused to an auxin-inducible degron^24,34^. We previously used this cell line to show that 3D loops and domains disappear after 6 hours of depletion of cohesin, and found that 146 genes change by more than 1.75-fold in expression levels^24^. Here, we applied the ABC model to build predicted maps of enhancer-gene regulatory interactions in HCT116 cells in the presence and absence of cohesin. To do so, we analyzed data collected from HCT116-RAD21-Degron cells treated or untreated with auxin for 6 hours^24^, using DNase-seq and H3K27ac ChIP-seq to estimate enhancer activity and ENCODE Hi-C data to estimate 3D contact frequency.

To understand the effects of cohesin depletion on enhancer-promoter physical and regulatory interactions, we first summarized the changes in 3D contact frequency between promoters and ABC predicted enhancers as measured by new, higher-resolution ENCODE Hi-C data^35^. Removing cohesin leads to an overall drop in contact frequency between promoters and ABC-predicted distal enhancers— with stronger decreases in 3D contact for element-promoter pairs located at longer distances (**Fig. 1a**). For example, element-promoter pairs located over 50Kb apart decreased in contact frequency by 65% on average, whereas pairs located less than 50Kb apart increased in contact frequency by 5% on average (**Fig. 1b**). This is consistent with a model where cohesin-mediated loop extrusion promotes the formation of longer-range 3D interactions, but does not have a strong impact on the frequency of shorter-range interactions, where polymer dynamics appear to be sufficient for frequent 3D contacts^36,37^. Notably, the reduction in 3D contacts for distal enhancers did not reach a baseline predicted by a simple power-law function of genomic distance (**Extended Data Fig. 1a**), suggesting that there may be additional mechanisms, such as compartmentalization effects, that maintain enhancer-promoter contacts even upon removal of cohesin^29–31^.

**Figure 1.**
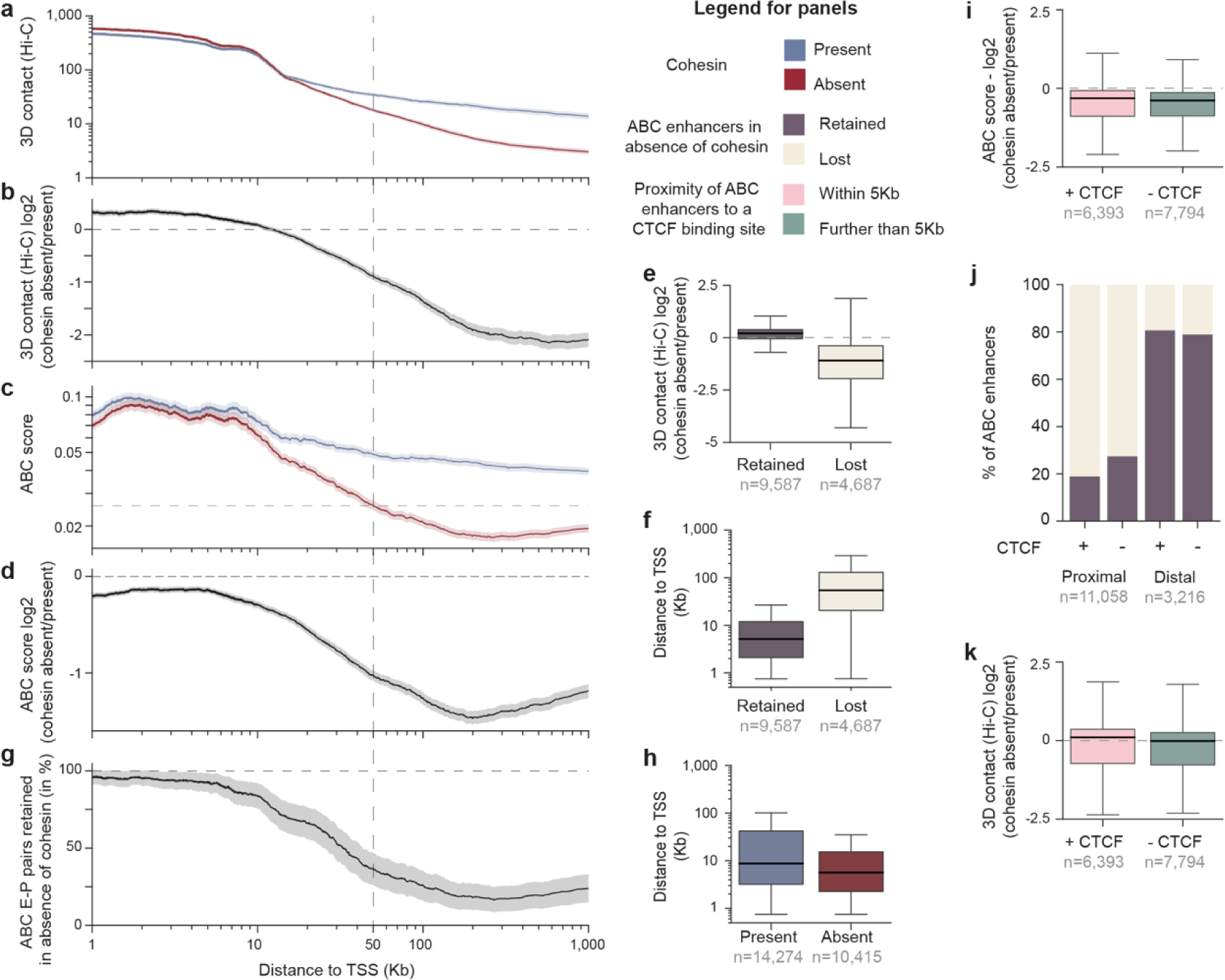
ABC predicts changes in enhancer effects upon cohesin removal. **a**, Enhancer-promoter 3D contact frequency (Hi-C SCALE-normalized counts, 5-Kb resolution^35^) in the presence (blue, no auxin, Hi-C depth = 1.1 billion contacts) or absence (red, plus 6 hours auxin, Hi-C depth = 1.5 billion contacts) of cohesin, as a function of distance from enhancer to target promoter. Lines: Rolling average for all enhancer-gene regulatory interactions predicted by ABC in the cohesin present condition (n=14,274) smoothed over 1,000 data point windows. Shading: 95% confidence interval (c.i.) of the rolling average. **b,** Similar to **a**, showing a log_2_ fold-change in Hi-C contact frequency between absence and presence of cohesin as a function of distance. Black line: Rolling average. Gray shading: 95% c.i. of the rolling average. **c,** Similar to **a**, showing ABC score as a function of distance. Dashed line represents the ABC score threshold (0.0257). **d,** Similar to **a**, showing a log_2_ fold-change in ABC score between the absence and presence of cohesin as a function of distance. **e,** Changes in 3D contact frequency (Hi-C SCALE-normalized counts, 5-Kb resolution, Hi-C depth = 1.1 (no auxin) and 1.5 (plus 6 hours auxin) billion contacts^35^) between cohesin present and absent, for all enhancer-promoter regulatory interactions predicted by ABC to be retained (dark gray, n=9,587) or lost (cream, n=10,415) in the absence of cohesin. Boxes: median and interquartile range. Whiskers: full distribution, except for points determined “outliers” using the interquartile range. **f,** Distribution of distances from enhancer to target promoter for all enhancer-promoter regulatory interactions retained (dark gray, n=9,587) or lost (cream, n=10,415) in the absence of cohesin. **g,** Similar to **a**, showing percentage of enhancer-promoter regulatory interactions predicted by ABC in the cohesin present condition that also pass the ABC score threshold (0.0257) in the absence of cohesin (retained enhancers, n=9,587), as a function of distance. **h,** Distribution of distances from enhancer to target promoter for all enhancer-promoter regulatory interactions predicted by ABC in the presence or absence of cohesin. **i,** Fold-changes in ABC scores between cohesin present and absent conditions, for all enhancer-promoter regulatory interactions near CTCF sites (+ CTCF; rose, n=6,393) or not near CTCF sites (-CTCF, sage, n=7,794) from CTCF ChIP-seq in HCT116. **j,** Percentage of enhancer-promoter regulatory interactions of distal and proximal ABC enhancers near or not near CTCF sites that are retained (dark gray) or lost (cream) in the absence of cohesin. **k,** Log_2_ fold-change in 3D contact frequency (Hi-C SCALE-normalized counts at 5-Kb resolution, Hi-C depth = 1.1 (no auxin) and 1.5 (plus 6 hours auxin) billion contacts^35^) between cohesin present and absent conditions, for all enhancer-promoter regulatory interactions near CTCF sites (+CTCF, rose, n=6,393) and not near CTCF sites (-CTCF, sage, n=7,794).

In subsequent analyses, we define elements as candidate enhancers based on DNase I hypersensitive sites (DHS) peaks and “enhancers” as a non-promoter element that is either predicted (by ABC) or validated to regulate a target gene. We further define “distal” elements as >50Kb, and “proximal” elements as <50Kb, based on the observation that the average change in contact frequency at 50Kb is ∼2-fold (**Fig. 1b**).

To explore how these changes in 3D contacts might affect enhancer-promoter regulatory interactions, we compared ABC predictions in the presence and absence of cohesin (**Supplementary Table 1**). We note that the ABC model assumes that the effect of an enhancer on target gene expression is proportional to the product of intrinsic enhancer activity and enhancer-promoter 3D contact frequency. Overall, enhancer activities between the two conditions were similar, whereas 3D contacts changed more dramatically (**Extended Data Fig. 1b**). As such, the fold-change in ABC scores between the presence and absence of cohesin are approximately equal to the fold-change in enhancer-promoter 3D contacts between the two conditions (**Extended Data Fig. 1c**).

In the presence of cohesin, ABC predicted 14,274 enhancer-gene regulatory interactions, including at least one ABC enhancer for 5,850 of 11,559 genes expressed at TPM > 1. In contrast, in the absence of cohesin, ABC predicted only 10,415 enhancer-gene regulatory interactions, with at least one ABC enhancer for only 5,190 genes (**Extended Data Fig. 1d-f**).

Similar to the changes in 3D contacts, ABC scores showed stronger decreases for element-promoter pairs located at longer distances, such that many no longer passed the threshold on the ABC score used to call a significant regulatory interaction (**Fig. 1c,d,e**). For example, 76% of enhancer-promoter pairs located over 50Kb apart were lost upon depletion of cohesin, compared to only 18% of enhancer-promoter pairs located less than 50Kb apart (**Fig. 1f,g**). As a result, the overall set of enhancer-promoter pairs predicted by ABC in the presence of cohesin included pairs spanning longer distances (90th percentile: 136Kb in presence of cohesin, versus 77Kb in the absence of cohesin, **Fig. 1h**).

ABC scores decreased not only for elements located near CTCF sites (within 5Kb of a CTCF ChIP-seq peak), but also for elements not located near CTCF sites (**Fig. 1i,j**). Apparently, 3D contacts decrease not only at CTCF-mediated loops, stripes, and domain boundaries, but also for element-promoter pairs located within contact domains (**Fig. 1k**).

We replicated these analyses with the newer and more accurate ENCODE-rE2G model^38^, which predicted the same relationship where distal enhancers are preferentially lost in the absence of cohesin (**Extended Data Fig. 2, Supplementary Table 1**).

Together, these analyses show that, in addition to the loss of loops and contact domains^24^, loss of cohesin leads to a strong decrease in quantitative contact frequency for candidate enhancer-promoter pairs located at long distances (*e.g.*, average decrease of 65% for pairs located >50Kb). Accordingly, the ABC model predicts that distal enhancers are strongly dependent on cohesin for their regulatory effects on gene expression, whereas enhancers located close to their target genes are not. Such an effect would be consistent with recent observations that recruiting synthetic transcription factors to either distal or proximal locations showed differential cohesin sensitivity^17^. We next set out to test this directly for endogenous enhancers in the genome using CRISPR perturbations.

### CRUDO measures the effect of 3D contacts on enhancer-promoter regulation

To experimentally test the role of 3D contact frequency in enhancer regulation, we developed CRISPRi of Regulatory elements Upon Degron Operation (CRUDO) — a highly sensitive method to test, for many enhancer-promoter pairs in parallel, whether their regulatory interactions depend on a particular protein of interest. In this approach, we apply CRISPRi pooled screens in a degron-engineered cell line to perturb putative enhancers in their endogenous context in the genome and measure the effects on gene expression in the presence and absence of the tagged protein (**Fig. 2a**). To read out effects on gene expression, we implemented CRUDO with either RNA fluorescence *in situ* hybridization and cell sorting (FlowFISH^18^), or single-cell targeted Perturb-seq (TAP-seq^39^) assays (see below, **Supplementary Table 2**).

**Figure 2.**
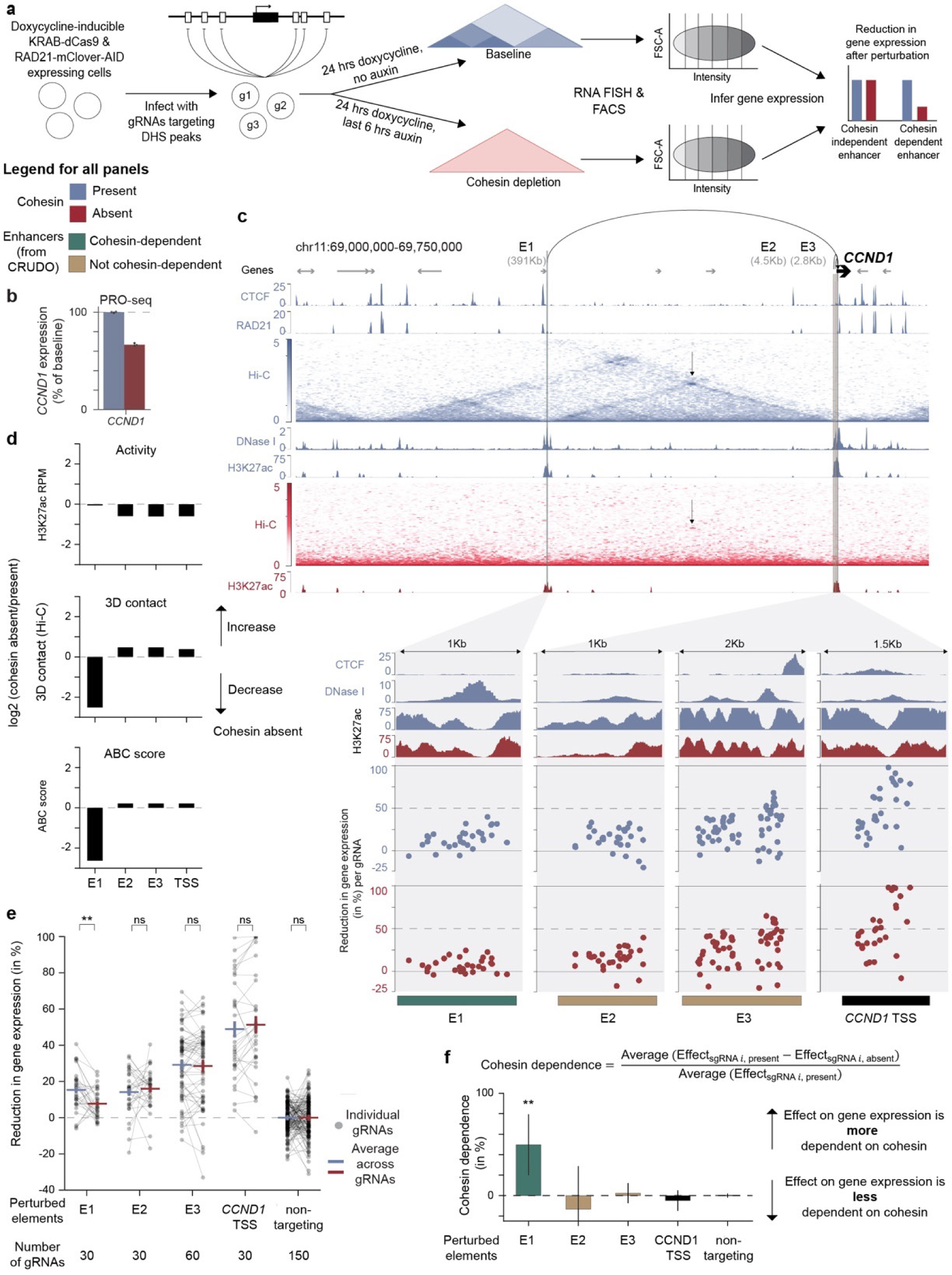
CRUDO-FF measures endogenous enhancer effects under controlled changes in quantitative contact frequency. **a**, Simplified schematic of the CRUDO-FF method for identifying the effects of enhancer-promoter contact frequency on enhancer effects. We start with cells expressing RAD21 tagged on both alleles with mClover and an auxin-inducible degron (AID) and dCas9-KRAB from a doxycycline-inducible promoter. Cells are transduced with a pool of gRNAs targeting DHS peaks surrounding the gene of interest. For the baseline condition (cohesin present, blue), cells are treated with doxycycline for 24h to activate the CRISPRi machinery. For the cohesin depletion condition (cohesin absent, red) the cells are treated with doxycycline for 24h to activate the CRISPRi machinery and in addition for the last 6h of the treatment auxin is added to degrade RAD21. Cells are then labeled for the gene of interest using RNA FISH and sorted based on their fluorescence levels into 6 different bins. gRNAs within each bin are sequenced, and their frequencies are analyzed to determine the quantitative effect of each gRNA on gene expression in the presence and absence of cohesin. Barplot shows exemplary results for a cohesin-independent enhancer that has the same effect on gene expression in the presence and absence of cohesin (blue and red bars are the same height) and a cohesin-dependent enhancer that has a stronger effect on gene expression in the presence compared to the absence of cohesin (blue bar is higher than red bar) indicating a higher dependence on cohesin for its regulatory effect. **b,** *CCND1* transcript levels in the presence (blue, 6h of DMSO treatment) and absence (red, 6h of auxin treatment) of cohesin as measured by PRO-seq. Error bars represent the 95% c.i. across 3 biological replicates. **c,** Browser snapshot of the *CCND1* locus showing DNase I hypersensitivity, CTCF ChIP-seq, RAD21 ChIP-seq, Hi-C at 5-Kb resolution (SCALE-normalized counts, Hi-C depth = 1.1 (no auxin) and 1.5 (plus 6 hours auxin) billion contacts^35^), and H3K27ac ChIP-seq in the presence (blue) and/or absence (red) of cohesin. Genomic coordinates in hg38: chr7:121,100,000-121,800,000. Zoom-ins: Bars at bottom show genomic locations of elements targeted in the pilot CRUDO-FF screen (enhancers E1-3 and the CCND1 TSS). Dots show the CRUDO-FF measured effects on *CCND1* expression for individual gRNAs targeting each element (% reduction in gene expression, normalized to the mean of all non-targeting gRNAs within each condition). **d,** Log_2_ fold-change in H3K27ac levels (top panel), 3D contact frequency (Hi-C SCALE-normalized counts at 5-Kb resolution, Hi-C depth = 1.1 (no auxin) and 1.5 (plus 6 hours auxin) billion contacts^35^, middle panel), and ABC score (bottom panel) in the absence versus presence of cohesin. **e,** CRUDO-FF measured effect sizes on *CCND1* expression for individual gRNAs in the presence (blue) or absence of cohesin (red). Horizontal lines and error bars: mean (+/-95% c.i.) of effect sizes for each element. ns: *P_BenjaminiHochberg_* > 0.05, **: *P_BenjaminiHochberg_* < 0.01 (gRNA-based paired 2-sample t-test). **f,** The level of cohesin dependence for each element, describing the change in the effect of an element on gene expression between the cohesin present versus absent conditions. Bar: mean across gRNAs for each element. Error bars: 95% c.i.. **: *P_BenjaminiHochberg_* < 0.01 (gRNA-based paired 2-sample t-test).

To apply CRUDO to study the effects of cohesin-mediated 3D contacts, we further engineered the previously generated HCT-116 RAD21 degron cell line^24,34^ to additionally express a dCas9-KRAB fusion (CRISPR interference, CRISPRi) linked to a blue fluorescent protein (BFP) under a doxycycline-inducible promoter. We confirmed that this doubly engineered cell line retained the previously observed gene expression changes upon depletion of cohesin (**Extended Data Fig. 3a**), and confirmed the efficacy of the dCas9-KRAB repression using previously validated guide RNAs (gRNAs, **Extended Data Fig. 3b,c, Supplementary Table 3**). We conducted CRUDO screens in this cell line by first transducing cells with a lentiviral library of gRNAs such that each cell received only one gRNA. We next induced dCas9-KRAB mediated enhancer repression by adding doxycycline for 24 hours and, in a second condition, we additionally depleted cohesin by adding auxin for the last 6 hours of the enhancer perturbation. This allowed us to measure changes in the strength of regulatory effects in the presence and absence of cohesin according to simulations considering mRNA half-lives and the rate of induced repression (see Methods, **Extended Data Fig. 4**). We harvested cells from each condition and measured the effects on gene expression using either FlowFISH or TAP-seq (see Methods). For FlowFISH screens, we used RNA FISH to label the mRNA of interest, sorted cells into six bins ranging from low to high target gene expression, used next-generation sequencing to measure the frequency of each gRNA in each bin, and used maximum likelihood estimation to infer the effects of each gRNA on gene expression compared to negative control (non-targeting) gRNAs in the same pool of cells (**Fig. 2a**). A key advantage of this CRUDO-FlowFISH (CRUDO-FF) approach is that we obtain precise measurements of the effect size of each candidate element on gene expression, by examining many thousands of single cells per gRNA, multiple gRNAs (2-164) per element, and hundreds to thousands of negative control gRNAs in a single pooled experiment.

To validate our approach, we first conducted a small CRUDO experiment to examine candidate enhancers for *CCND1* — a cohesin-sensitive gene whose transcription decreases by 33.4% upon depletion of cohesin, as measured by precision run-on sequencing (PRO-seq, **Fig. 2b**). We designed 30 gRNAs each for the promoter of CCND1 and three of the ten predicted ABC enhancers. All elements showed comparable levels of H3K27ac in the presence and absence of cohesin, indicating the elements do not change in their intrinsic activity (**Fig. 2c,d**). While enhancer E1 showed a decrease in 3D contact with the *CCND1* promoter in the absence of cohesin, the other two enhancers did not **(Fig. 2d,e)**, resulting in only E1 being predicted by ABC to decrease in its effect size **(Fig. 2d).** We performed CRUDO-FF, and found that perturbations to all 3 targeted enhancers significantly reduced gene expression relative to negative control gRNAs both in the presence and absence of cohesin (**Fig. 2c,e)**.

We next calculated the degree to which the quantitative effect of each enhancer-gene regulatory interaction was dependent on cohesin. We defined “cohesin dependence” as the degree to which the strength of the regulatory effect of an enhancer-gene pair decreases upon removal of cohesin, expressed as a percentage of the effect in the presence of cohesin (**Fig. 2f**).

For example, if perturbation to an enhancer leads to a 50% decrease in gene expression in the presence of cohesin and a 0% decrease in gene expression in the absence of cohesin, its effect is fully dependent on cohesin (“cohesin dependence” = 100%). In contrast, if perturbation to an enhancer leads to a 50% decrease in gene expression in both conditions, its effect is independent of cohesin (cohesin dependence = 0%). We note that this calculation assumes that enhancers should have a consistent relative effect on gene expression across the two conditions (that is, enhancer activity combines multiplicatively with the promoter to determine gene expression), as previously observed in reporter assays^40,41^. We tested for statistical significance by assessing the consistency of the effects across all gRNAs targeting a given element, and defined significantly “cohesin dependent” pairs based on a two-sample paired t-test (*P_BenjaminiHochberg_* < 0.05).

In this *CCND1* pilot experiment, the promoter and two proximal enhancers were not cohesin dependent: they showed similar decreases in gene expression between the cohesin present and absent conditions (promoter: 48.8% and 51.2%; enhancers: 14.1% and 16%, 29.1% and 28.4%, cohesin present and absent, respectively; cohesin dependence = -4.9%, -13.3%, and 2.3%, respectively, **Fig. 2c,e,f**). In contrast, the third enhancer (E1) had a significantly smaller effect on gene expression in the absence of cohesin (cohesin present: 15.3%, cohesin absent: 7.7%, *P_BenjaminiHochberg_* = 0.009; cohesin dependence = 49.7% (**Fig. 2c,e,f**)). Notably, these effects were consistent with the ABC predictions (**Fig. 2d)**, because only E1 changes in its contact frequency with the *CCND1* promoter in the absence of cohesin (**Fig. 2c,d**). Thus, of the three tested distal enhancers, one (E1) appears cohesin dependent and the others (E2 and E3) not (**Fig. 2i)**.

These data demonstrate our ability to measure perturbation-induced changes in gene expression both in the presence and absence of cohesin and identify cases of cohesin-dependent and - independent enhancer-gene regulatory interactions. Thus, CRUDO provides an approach to quantitatively measure the effect of enhancer-promoter contact frequency on enhancer function in the native genomic context.

### Distal enhancer-gene regulatory interactions depend on cohesin

We next applied CRUDO to comprehensively characterize all enhancers for five representative cohesin-sensitive genes — CCND1, FAM3C, KITLG, SSFA2, and MYC — selected on the basis of their high expression levels (enabling good detection) and the presence of ABC-predicted enhancers that exhibited diverse changes in 3D contact frequency in the absence of cohesin (**Fig. 3, Extended Data Fig. 5, 6, 7, 8**). For each gene, we used CRUDO to target each element within 1Mb and measure effects on gene expression using FlowFISH in the presence or absence of cohesin (FlowFISH probesets are listed in **Supplementary Table 4**).

**Figure 3:**
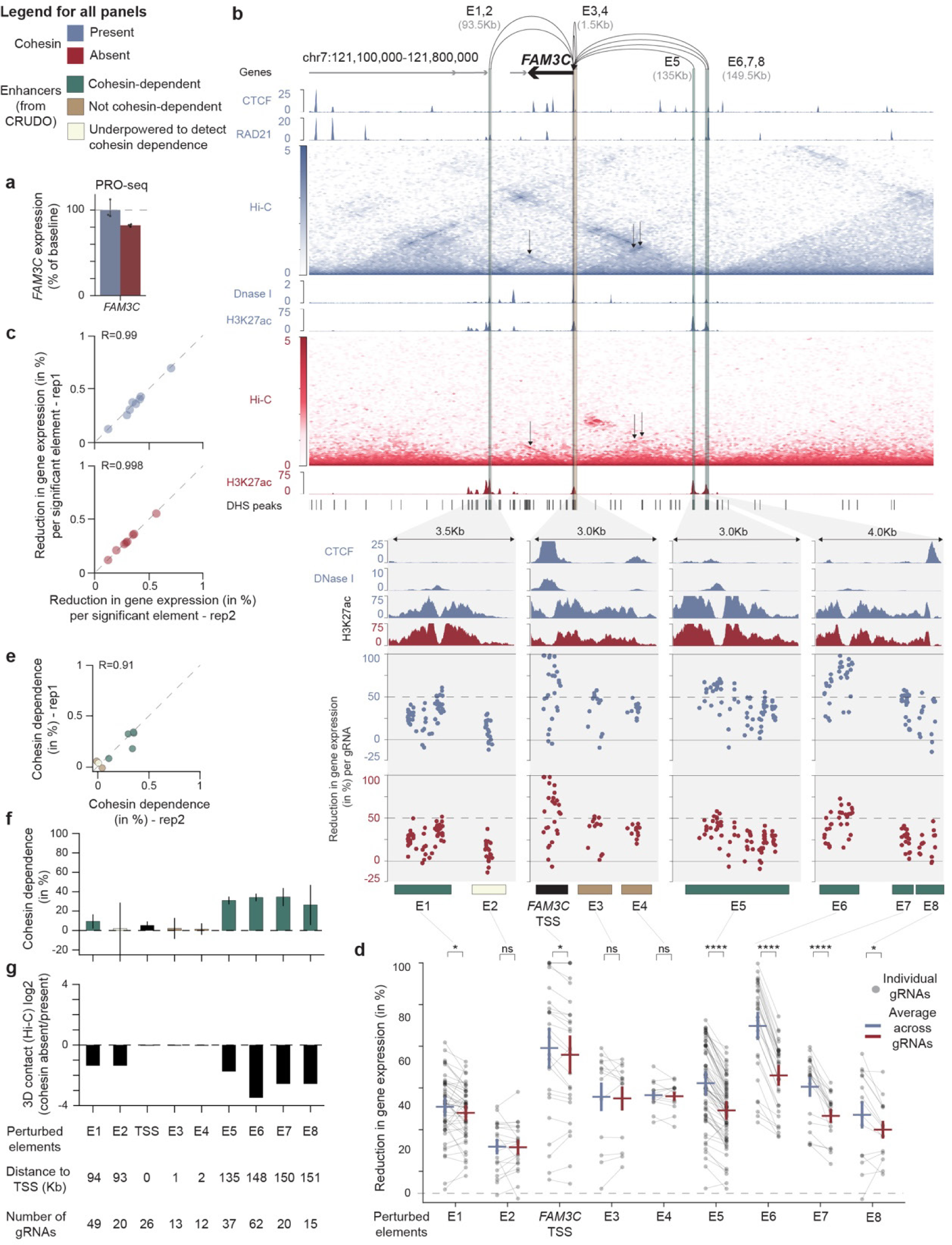
Regulatory interactions of distal enhancers decrease in the absence of cohesin. **a**, *FAM3C* transcript levels in the presence (blue, 6h of DMSO treatment) or absence (red, 6h of auxin treatment) of cohesin, as measured by PRO-seq. Error bars: 95% c.i. across 3 biological replicates. **b,** *FAM3C* locus showing DNase I hypersensitivity, CTCF ChIP-seq, RAD21 ChIP-seq, Hi-C at 5-Kb resolution (SCALE-normalized counts, Hi-C depth = 1.1 (no auxin) and 1.5 (plus 6 hours auxin) billion contacts^35^), and H3K27ac ChIP-seq in the presence (blue) and/or absence (red) of cohesin. Bars at bottom show locations of all elements tested in the CRUDO-FF experiment. Zoom-ins show all elements that had a significant effect on *FAM3C* expression in cohesin present condition. Bars at bottom show genomic locations of elements identified in the CRUDO-FF screen (teal: cohesin-dependent enhancers; taupe: not cohesin-dependent enhancers; ivory: underpowered to detect cohesin dependence enhancers; black: TSS). Dots show the measured effects on *FAM3C* expression for individual gRNAs targeting each element (% reduction in gene expression, normalized to the mean of all non-targeting gRNAs within each condition). **c**, Scatterplot shows correlation of effects on gene expression between two biological replicate experiments in the cohesin present (blue, top panel) and cohesin absent (red, bottom panel) conditions. Dots: all 9 elements that had significant effects on gene expression in the cohesin present condition. **d,** CRUDO-FF measured effect sizes on *FAM3C* expression for individual gRNAs in the presence (blue) or absence of cohesin (red). Horizontal lines and error bars: mean (+/- 95% c.i.) of effect sizes for each element. ns: *P_BenjaminiHochberg_* > 0.05, *: *P_BenjaminiHochberg_* < 0.05, ****: *P_BenjaminiHochberg_* < 0.0001 (gRNA-based paired 2-sample t-test). **e,** Similar to panel **c,** showing correlation of cohesin dependence between two biological replicate experiments. **f**, The level of cohesin dependence for each element, describing the change in the effect of an element on gene expression between the cohesin present versus absent conditions. Bar: mean across gRNAs for each element. Error bars: 95% c.i.. **g**, Log_2_ fold-change in 3D contact frequencies (Hi-C SCALE-normalized counts at 5-Kb resolution, Hi-C depth = 1.1 (no auxin) and 1.5 (plus 6 hours auxin) billion contacts^35^) between the absence and presence of cohesin.

For example, we examined *FAM3C*, a gene whose transcription decreases by 17.9% upon depletion of cohesin (**Fig. 3a**). Of 96 tested candidate enhancers in the locus, we identified 8 that significantly affected the expression of *FAM3C* in the baseline (cohesin present) condition, located between 1-151Kb from the FAM3C promoter (**Fig. 3b**). The effect sizes of these 8 enhancers ranged from 12-70%, and were highly correlated across two biological replicate experiments (Pearson’s *R* = 0.99, **Fig. 3c)**. 5 of the 8 enhancers had effects on *FAM3C* expression that were significantly cohesin-dependent (**Fig. 3b,d,e,f**). On average, these 5 cohesin-dependent enhancers showed a 73% decrease in enhancer-promoter contact frequency in the absence of cohesin, compared to only 7% for the enhancers that were not significantly cohesin-dependent (**Fig. 3g**). Interestingly, the cohesin-dependent enhancers were all located at least 94Kb from the FAM3C promoter, while the other enhancers varied in genomic distance (1–93Kb, **Fig. 3b,f,g**).

Considering all five genes, we tested a total of 1,039 putative element-gene pairs (96-575 per gene), and identified 34 significant element-gene pairs in the presence of cohesin (4-10 per gene) (**Fig. 4a**). Of these 34, 26 showed a nominal (>5%) decrease in regulatory effects in the absence of cohesin, whereas 8 did not. After significance testing, we called 18 element-gene pairs as “cohesin-dependent” (significant decrease in their regulatory effects in the absence of cohesin, *P*_Benjamini-Hochberg_ < 0.05), 8 as “not cohesin-dependent” (no significant decrease, despite having >90% power to detect 50% cohesin dependence or a significant increase in their regulatory effects in the absence of cohesin), and 8 as “underpowered to detect cohesin dependence” (<90% power to detect 50% cohesin dependence) (**Fig. 4a**, **Extended Data Fig. 10a,b**).

**Figure 4:**
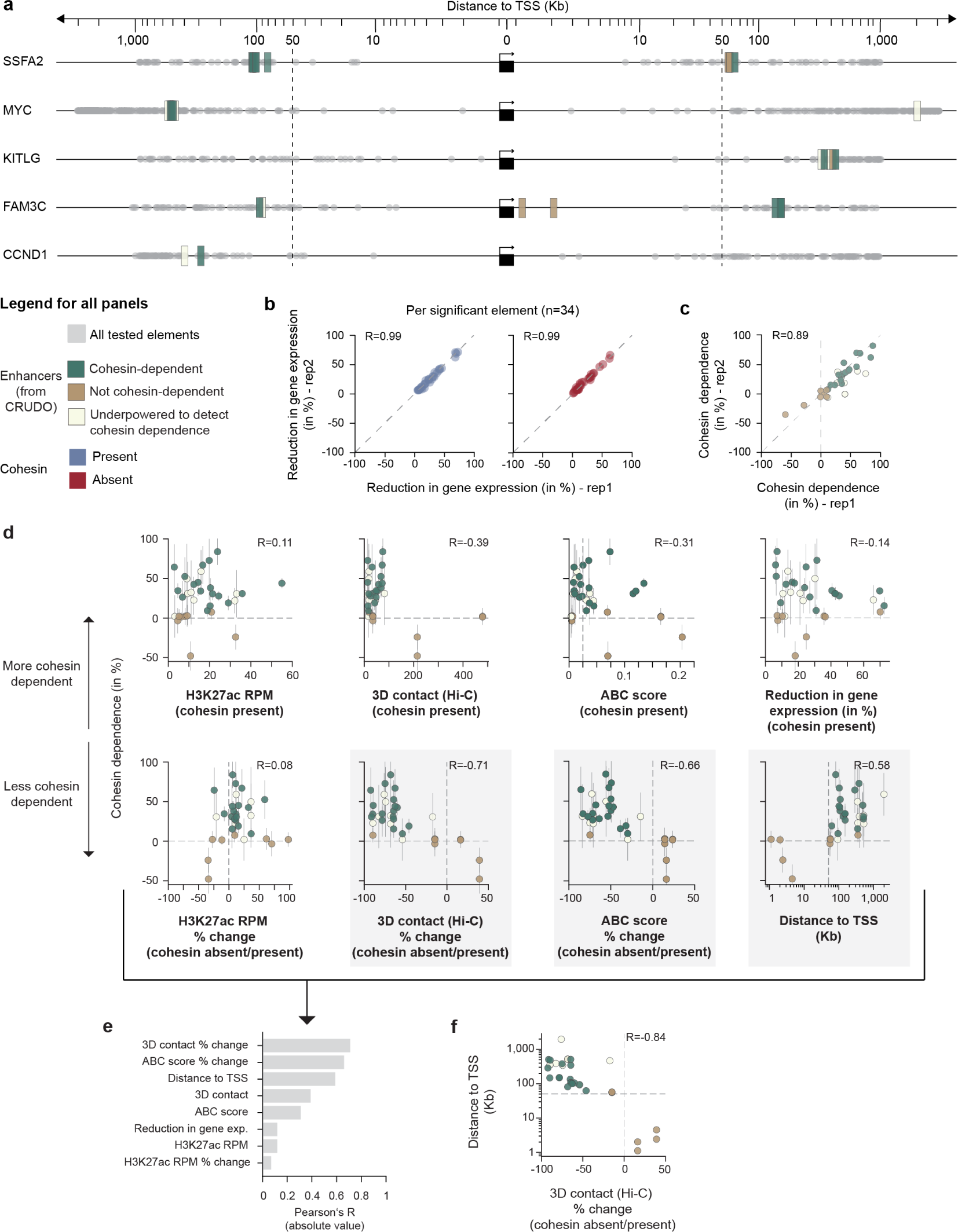
Enhancer-promoter contact frequency modulates enhancer effects. **a**, Summary of enhancers identified by CRUDO-FF across 5 genes, showing the locations of enhancers (34 total) and their cohesin dependence relative to the target gene TSS. Teal: Cohesin-dependent enhancers (n=18). Taupe: Not cohesin-dependent enhancers (n=8). Ivory: Underpowered to detect cohesin dependence enhancers (n=8). Light gray: All tested elements (n=1,039). **b**, Scatterplot shows correlation of element-level effects between two biological replicate experiments in the cohesin present (blue) and absent (red) conditions. **c,** Scatterplot shows correlation of cohesin dependence as estimated from two separate biological replicate experiments. **d,** Comparison of cohesin dependence with other genomic features across all enhancers (n=34). Top from left to right: H3K27ac ChIP-seq signal at the enhancer in the cohesin present condition; 3D contact (SCALE-normalized counts, 5-Kb resolution, Hi-C depth = 1.1 billion contacts^35^) in the cohesin present condition; ABC-score in the cohesin present condition; Reduction in gene expression in the cohesin present condition. Bottom from left to right: Percent change in H3K27ac ChIP-seq signal at the enhancer between the cohesin absent versus present conditions; Percent change in 3D contacts between the cohesin absent versus present conditions; Percent change in ABC score between the cohesin absent versus present conditions; Genomic distance between the enhancer and target gene TSS in Kb (log_10_ scale). Error bars: 95% c.i. across individual gRNAs (n=10-69). **e**, A bar graph illustrating Pearson’s R values, derived from scatter plots in **d**, indicating the correlation between cohesin dependence and genomic features: percent change in H3K27ac ChIP-seq signal between cohesin absent versus present conditions, H3K27ac ChIP-seq signal, reduction in gene expression, ABC score, 3D contact (Hi-C SCALE-normalized counts, 5-Kb resolution, Hi-C depth = 1.1 billion contacts^35^), genomic distance between the enhancer and target gene TSS in Kb (log_10_ scale), percent change in ABC score between cohesin absent versus present conditions, and percent change in 3D contacts between cohesin absent versus present conditions across all enhancers (n=34). **f,** Genomic distance (log_10_ scale) from enhancer to target gene TSS versus the percent change in 3D contacts (Hi-C SCALE-normalized counts, 5-Kb resolution, Hi-C depth = 1.1 (no auxin) and 1.5 (plus 6 hours auxin) billion contacts^35^) between the absence and presence of cohesin.

The effect sizes of the 34 regulatory element-gene pairs were highly correlated across biological replicate experiments (Pearson’s R = 0.99 for both cohesin present and absent conditions, respectively, **Fig. 4b**), as was the degree of cohesin dependence (Pearson’s R = 0.89, **Fig. 4c**). To validate these effects, we tested 24 elements in a second experiment using a different assay (CRUDO-TAP-seq, **Methods, Supplementary Table 2**), and found that enhancer effects and cohesin dependence were correlated between FlowFISH and TAP-seq (Pearson’s R = 0.65 for effects at baseline; Pearson’s R = 0.59 for cohesin dependence; **Extended Data Fig. 9**).

We characterized the relationship between cohesin dependence and 3D contacts across these 34 regulatory elements for 5 genes. Changes in enhancer effect on gene expression (cohesin dependence, as measured by CRUDO) were correlated with the change in enhancer-promoter 3D contact frequency (as measured by Hi-C) (Pearson’s R = –0.71, **Fig. 4d,e**), as well as with the change in ABC score (Pearson’s R = –0.66). This correlation held for elements regardless of their proximity to a CTCF binding site (**Extended Data Fig. 10c**), and was stronger than with other genomic features, such as baseline H3K27ac, baseline 3D contacts, baseline effect sizes, baseline ABC score, and changes in H3K27ac (absolute Pearson’s R = 0.08 – 0.39, **Fig. 4d,e**). Among the 18 enhancer-gene pairs that were significantly cohesin dependent, 17 showed a >2-fold decrease in 3D contact upon removal of cohesin, versus only one of 8 not cohesin-dependent enhancer-gene pairs (odds ratio: 75, Fisher exact *P* = 0.00009). Conversely, of the 25 enhancer-gene pairs that changed in 3D contact by more than 2-fold, 17 were significantly cohesin-dependent, versus one of the 8 pairs that changed in 3D contact by less than 2-fold.

We also find a correlation between cohesin dependence and the linear distance between the enhancer and its target promoter (Pearson’s R = 0.59, P = 0.00023, **Fig. 4d,e**). Strikingly, all of the cohesin-dependent enhancers were also located further than 50Kb from their target promoter (**Fig.4a,d**), consistent with our observations that 3D contact frequencies are more sensitive to cohesin above this distance (**Fig. 1a,b**). The effects of distal enhancers (>50 Kb) on gene expression were 6.6-fold more likely to decrease upon removal of cohesin, compared to very proximal enhancers (<10 Kb) (Fisher exact *P* = 0.0392). This suggests that distal enhancers are more sensitive to a reduction in effect upon the removal of cohesin (Pearson’s *R* = -0.84, P = 4.7 x 10^-10^, **Fig.4f**). Interestingly, it appears that these cohesin-sensitive genes are predominantly regulated by distal enhancers, with only a small fraction of enhancers being very proximal. Moreover, the absence of enhancers located within the 10-50 Kb range could suggest a potential contribution of the enhancer landscapes in modulating the sensitivity of genes to cohesin.

Together, we find that a decrease in enhancer-promoter contact frequency (>2-fold) typically coincides with a dependence on cohesin for the regulatory effect of that enhancer. These data demonstrate that changing 3D contact frequencies quantitatively tunes the regulatory effects of enhancers on gene expression. This effect is particularly pronounced at distal enhancer-promoter pairs (>50Kb).

### Cohesin-dependent genes are primarily regulated by distal enhancers

Our data indicate a quantitative relationship between enhancer-promoter contact frequency and enhancer effects on gene expression, and that distal enhancers rely on cohesin to mediate these interactions. Accordingly, we next investigated whether the sensitivity of genes to cohesin is related to features of their enhancer landscapes — specifically, to how many, how far away, and/or how strong a gene’s enhancers might be.

We compared the enhancers for the five cohesin-sensitive genes (as measured by CRISPRi-FlowFISH in the cohesin present condition) to a previous study in which we performed similar comprehensive tiling experiments on 23 “tiling control genes” that were not selected based on information about their sensitivity to cohesin depletion^18^. On average, the 5 cohesin-dependent genes we studied here were regulated by 6.8 enhancers, whereas the 23 tiling control genes had an average of 3.1 enhancers (**Extended Data Fig. 11a**). We observed distinct differences in the characteristics of the identified enhancers for cohesin-dependent genes compared to the control group: 88.2% (30/34) of the enhancers for cohesin-dependent genes were located distally (>50Kb), whereas only 23.9% (17/71) of enhancers were distal in the tiling control gene set (**Fig. 5a, Extended Data Fig. 11b,c**). Moreover, when analyzing the effect sizes of the distal enhancers, we found that none of the distal enhancers regulating genes in the tiling control gene set exhibited more than a 20% effect size, while more than half (16/30) of the distal enhancers regulating cohesin-dependent genes had effects exceeding 20% (**Fig. 5b, Extended Data Fig. 11b,c**). These findings indicate that cohesin-dependent genes tend to be regulated by more enhancers, and that those enhancers are more distal with stronger effect sizes.

**Fig 5:**
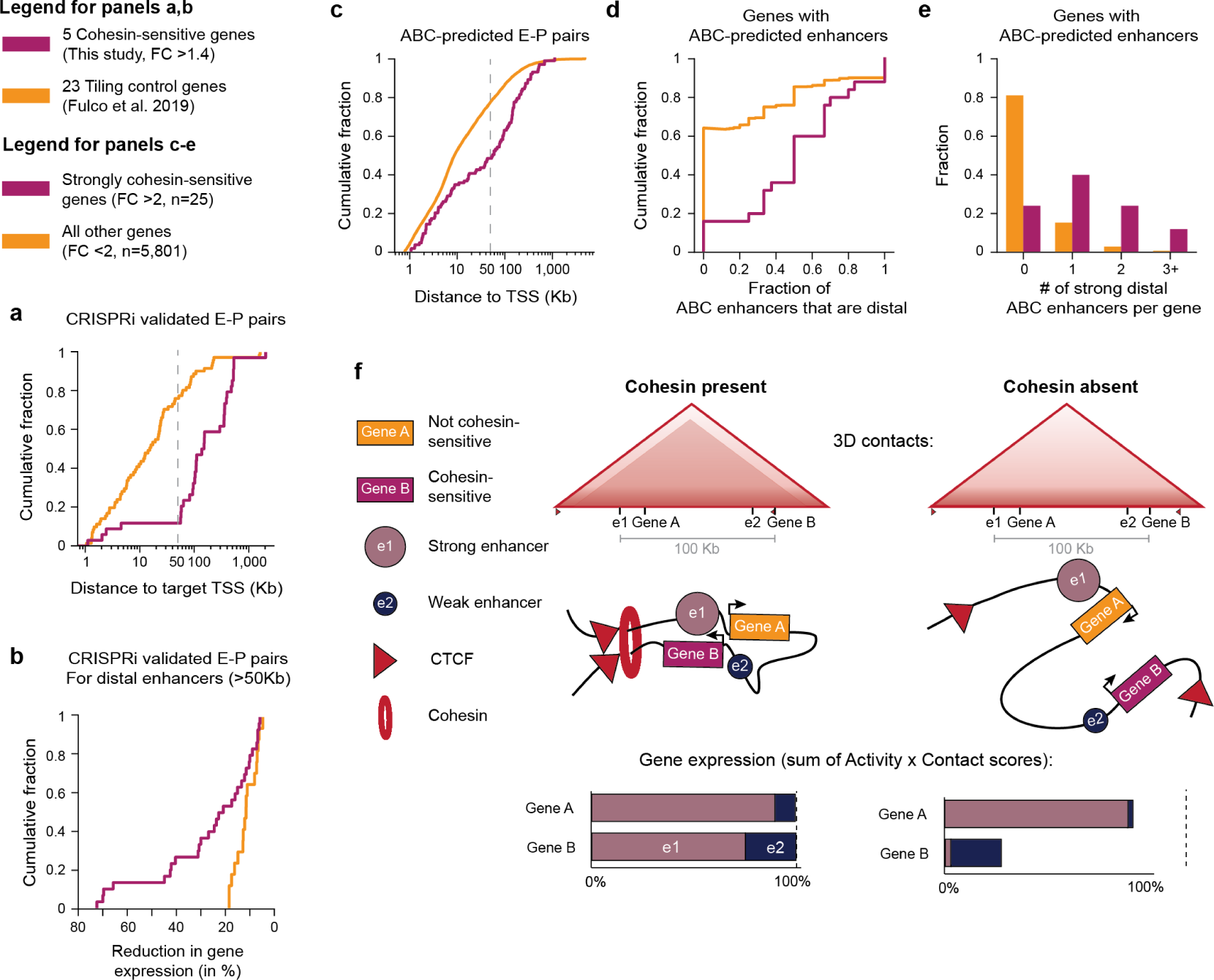
Cohesin-dependent genes rely on distal enhancers. **a,** Cumulative distribution plot of the genomic distance from enhancer to target gene TSS for each significant pair from comprehensive CRISPRi tiling data for the 5 cohesin-dependent genes in HCT116 cells (purple, n=34 this study) and for the 23 “Tiling control” genes we previously studied in K562 cells, which were selected without respect to their cohesin dependence (peach, n=71; Fulco et al.^18^). **b,** Cumulative distribution plot of the effect size for each distal enhancer (>50Kb distance to TSS) in the CRISPRi validated enhancer-promoter pairs from our CRUDO-FF dataset (purple, n=30) and the “Tiling control” dataset^18^ (peach, n=71).**c,** Cumulative distribution plot of the distance from enhancer to target gene TSS for each ABC-predicted enhancer-promoter pair in HCT116 cells for strongly cohesin-dependent genes (purple, n=103; defined as a >2-fold reduction in PRO-seq data in cohesin absent versus present conditions, TPM > 1) and all other genes (peach, n=14,171, TPM > 1). **d,** Considering genes that have at least one predicted ABC enhancer, cumulative distribution plot of the fraction of predicted enhancers that are distal (>50Kb distance to TSS) for strongly cohesin-dependent genes (purple, n=25) and all other genes (peach, n=5,803). **e,** Considering genes that have at least one predicted ABC enhancer, fraction of strongly cohesin-dependent genes (purple, n=25) and all other genes (peach, n=5,803) that have 0, 1,2, or more than 3 ABC-predicted strong, distal enhancers (>20% effect size, >50Kb distance to TSS). **f,** Proposed model in which the frequency of enhancer-promoter physical interactions quantitatively tunes the effects of enhancers on gene expression and where the sensitivity of genes to cohesin depletion depends on the relative strengths and genomic locations of their enhancers. In the presence of cohesin (left), loop extrusion forms a contact domain bounded by two CTCF binding sites (top), which increases contact frequencies between e1, e2, Gene A, and Gene B (middle). In the Activity-by-Contact model, the expression of each gene would be explained by the sum of enhancer Activity x Contact scores (bottom). Upon removal of cohesin (right), 3D contacts between more distal pairs of enhancers and promoters (e1-Gene B and e2-Gene A) are reduced, whereas 3D contacts between more proximal pairs of enhancers and promoters (e1-Gene A and e2-Gene B) are maintained due stochastic chromatin interactions. Due to the differences in the activities and 3D contacts of their respective enhancers, the two genes show differential sensitivity to cohesin: Gene A is less sensitive because it has a strong proximal enhancer, whereas Gene B is more sensitive because its strong enhancer is distal. Our analyses indicate that, across the genome, the pattern shown by Gene B (strong distal enhancer) is rare (only 19% of genes).

Moreover, we observed similar results when comparing the 5 cohesin-dependent genes to a larger set of 240 control genes studied using CRISPRi enhancer perturbation experiments with alternative designs that did not involve comprehensive tiling around individual genes (“Other control genes”, ^39,42–50^, **Extended Data Fig. 11d,e**)

To determine whether these differences hold for genes beyond the five that we experimentally studied here, we compared properties of enhancers predicted by the ABC model between genes that are strongly cohesin dependent (>2-fold decrease in transcription upon depletion of cohesin, n=25) and genes that are not (<2-fold decrease, n=5,803). Cohesin-dependent genes indeed had enhancers located farther away than for other genes (75^th^ percentile: 152Kb vs 42 Kb, Mann-Whitney U-test *P* = 1.5 x 10^-7^, **Fig. 5c**). 64% of cohesin-dependent genes are predicted to be regulated by mostly (>50%) distal enhancers (compared to just 24% of other genes; odds ratio = 5.6, Fisher exact P = 0.000026, **Fig. 5d**), and 76% had at least one distal enhancer predicted to have a >15% effect size (compared to just 19% of other genes; odds ratio = 13.5, Fisher exact P =1.6 x 10^-9^, **Fig. 5e, Extended Data Fig. 12**).

Together, these observations show that cohesin-dependent genes have distinct enhancer landscapes: they tend to have more total enhancers, more distal enhancers, and more strong distal enhancers than other genes.

## Discussion

The role of 3D enhancer-promoter contacts in gene regulation is the topic of ongoing debate^51–53^. The needed experiment — to manipulate the 3D contacts of enhancers in their endogenous locations in the genome to accurately measure effects on enhancer-promoter regulation — has proven challenging, and has been attempted at only a handful of enhancers, with mixed and subtle effects^9,13–17^. Here, we develop CRUDO to systematically test the role of 3D contacts on gene regulation at thousands of candidate enhancer-promoter pairs—producing the largest dataset to date describing the responses of many enhancer-promoter regulatory interactions to changes in 3D contacts in their endogenous genomic locations.

Our data demonstrate that reducing 3D contacts between an enhancer and target promoter almost always leads to a detectable reduction in enhancer effects on gene expression. Specifically, of the 25 regulatory enhancer-promoter pairs with reductions in 3D contact upon removal of cohesin (by >50%, **Fig. 4d**), 17 element-gene pairs were classified as cohesin-dependent” (17/25, 68%), 1 as “not cohesin-dependent” (1/25, 4%), and 7 as “underpowered to detect cohesin dependence” (7/25, 28%, **Fig. 4d**). These changes in regulatory effects were often small in magnitude (5-24% relative to baseline gene expression), and were only possible to detect due to the high sensitivity of our CRUDO-FF approach.

Together with previous observations, our results suggest a model for how cohesin-mediated 3D contacts tune enhancer function. Cohesin controls 3D genome organization through loop extrusion^19,21,54^. Upon removal of cohesin, CTCF-mediated loops and domains disappear, and 3D contacts change across the genome^24,25^. However, the magnitude of these changes depends on genomic distance. At shorter distances, 3D contacts are largely maintained (<5% changes, **Fig.1a,b**), presumably by more random, non-extrusion movement of the DNA polymer. At longer distances, 3D contacts between certain enhancers and promoters often change more dramatically (*e.g.*, by up to 89.83% **Fig. 3g**), for example, due to the interplay between cohesin and CTCF at domain boundaries^8,13,55,56^. Our data show that these differences in 3D contacts lead to quantitative changes in the effects of individual enhancers on their target genes — with changes in gene expression correlated with the change in 3D contact (**Fig. 4d**). As such, proximal enhancers (*e.g.*, 1-5 Kb) do not change appreciably in 3D contact frequency nor effect on gene expression, whereas more distal enhancers (*e.g.*, >50Kb) tend to show larger changes, consistent with previous studies of individual enhancer-promoter pairs in exogenous loci^11,12^.

Our data help to resolve questions as to why the removal of architectural proteins like cohesin or CTCF leads to only subtle changes in gene expression^23–25,33^ — such as, for example, only 146 genes changing by more than 1.75-fold upon removal of cohesin in HCT-116 cells^24^. Previous studies have attributed these subtle changes in gene expression to a variety of different factors, including proximity to super-enhancers^24,33^, CTCF binding at promoters^16^, presence of long-range loops^32^, a requirement for CTCF in establishing but not maintaining gene expression^31^, and the ability of enhancers and promoters to maintain a degree of focal contact even without cohesin due to compartmentalization effects^24^. Our results include three observations that together provide a refined explanation (not necessarily exclusive with some others). (i) Rather than a global requirement for cohesin at all enhancers, we find that cohesin quantitatively tunes 3D contacts and enhancer effects on gene expression, with much stronger effects for longer-range enhancers. (ii) Enhancers often have small effects on gene expression (median effect size in our combined CRISPR perturbation datasets = 10.9%^18^). (iii) Only a small number of genes have strong distal enhancers. Based on the ABC model and perturbation results, less than 20% of genes are predicted to have a strong, distal enhancer (>15% effect located >50Kb away, **Fig. 5e**). Together, due to the stronger influence of cohesin at distal enhancers and the distribution of enhancer locations in the genome, only a small number of genes would be predicted to have cohesin-dependent enhancers, and therefore the changes in gene expression due to acute removal of cohesin are expected to be small for most genes (**Fig. 5f**). Nevertheless, small changes to gene expression can have functional consequences, and we expect that many of these cohesin-dependent effects could have functional roles in the cell which then lead to further and larger downstream changes in gene expression, as previously proposed to occur during macrophage activation^57^. For example, Cyclin D1 (CCND1), one of the cohesin-dependent genes we study here, is a key cell cycle regulator involved in the G1 to S transition^58^. Reduced CCND1 expression has been shown to induce G0/1 arrest and a slower progression through S-phase^59^, while increased CCND1 expression —frequently observed in cancer — is associated with an accelerated transition through the cell cycle^60^, highlighting the importance of tightly controlled CCND1 expression levels.

While we found that quantitative changes in 3D contacts and regulatory effects of enhancers were correlated (Pearson’s R = –0.71, **Fig. 4d**), the relationship was far from perfect, and we were unable to address questions such as whether regulatory effects vary linearly^18,61^ or nonlinearly^11^ with respect to 3D contact. This may be due to other technical or biological factors that highlight directions for future work. First, while we have carefully controlled the timing of cohesin removal using an auxin-inducible degron, the temporal resolution of CRUDO-FF is still limited by the half-life of target mRNAs and induction time of dCas9-KRAB. Due to these factors, it is possible that our measurements of cohesin dependence underestimate the true effects (*i.e.*, due to mature mRNA levels not representing instantaneous effects on transcription) or conflate the effects of direct, *cis* effects with indirect, *trans* effects (*i.e.*, due to downstream effects of changing *cis* target gene expression during the 24 hours of CRISPRi repression). Second, estimates of 3D contact frequencies depend on the resolution at which Hi-C data is collected and analyzed^61,62^, and so collecting Hi-C data at even greater depth may improve our analysis. Third, we find that changing 3D contact frequencies, as measured by Hi-C, changes enhancer regulation, but these observations are possibly compatible with various models for defining functional enhancer “contacts”, including ones where enhancer and promoter DNA may be separated by dozens of nanometers or where functional “contacts” involve larger clusters of transcriptional factors and cofactors^27,57,63,64^. Finally, additional biological changes that occur with cohesin depletion could affect the relationship between pairwise enhancer-promoter contacts and their regulatory effects, including for example changes in enhancer-enhancer contacts that can impact regulation^38^ or changes in the sensitivity of promoters to distal enhancers^40,41^. Future work with additional temporally or spatially resolved assays will help to address these questions and further refine our understanding of the functional relationship between genome architecture and enhancer effects on gene expression.

In summary, we developed an approach to systematically investigate how cohesin-mediated 3D contacts influence gene expression, revealing general rules of gene regulation that will aid in studying how cohesin and distal enhancers coordinate cellular differentiation and signaling pathways in health and disease. Notably, our approach can be adapted to examine the role of other architectural or transcriptional regulatory proteins in enhancer regulation, which we expect will further our ability to differentiate between chromatin architecture as a packaging tool and its function in regulating gene expression.

## Supplementary tables

Supplementary Table 1: Enhancer-gene predictions for HCT116 cells

Supplementary Table 2: Datasets of experimentally tested noncoding element - gene pairs

Supplementary Table 3: gRNA designs for validation experiments

Supplementary Table 4: FlowFISH probe sets

Supplementary Table 5: qPCR primer sequences

## Data availability

CRISPRi-FlowFISH data: IGVF Data Portal (Accession numbers in Supplementary Table 2a)

CRISPRi-TAP-seq data: IGVF Data Portal (Accession numbers in Supplementary Table 2a)

ENCODE Hi-C from the ENCODE Portal: ENCSR958BEA, ENCSR087JOM

DNase-seq from the ENCODE Portal: ENCSR823GSV

ChIP-seq data from Gene Expression Omnibus: GSE104888

PRO-seq data from Gene Expression Omnibus: GSE106886

ENCODE-rE2G CRISPRi benchmarking set:

https://github.com/EngreitzLab/CRISPR_comparison/blob/main/resources/crispr_data/EPCrisprBenchmark_ensemble_data_GRCh38.tsv.gz.

## Code availability

ABC: https://github.com/broadinstitute/ABC-Enhancer-Gene-Prediction

ENCODE-rE2G: https://github.com/karbalayghareh/ENCODE-rE2G

FlowFISH processing pipeline: https://github.com/EngreitzLab/crispri-flowfish

TAP-seq primer design pipeline: https://github.com/argschwind/TAPseq

TAP-seq processing pipeline: https://github.com/argschwind/TAPseq_workflow

Code for additional analyses and visualization is available upon request and will be uploaded to a public repository prior to publication.

## Acknowledgements

P.G. was supported by a PhD fellowship from the Boehringer Ingelheim Fonds. B.R.D. was supported by the National Science Foundation Graduate Research Fellowship (DGE-1656518) and a Stanford Interdisciplinary Graduate Fellowship affiliated with Stanford Bio-X. Y.T. was supported by the National Science Foundation Graduate Research Fellowship (DGE-2146755). M.U.S. acknowledges the support of an NSF Graduate Research Fellowship (DGE-1656518) and was supported by a graduate fellowship award from Knight-Hennessy Scholars at Stanford University. A.R.G. and L.M.S acknowledge the support of R01HG011664 and the NHGRI Impact of Genomic Variation on Function Consortium (UM1HG011972). M.T.K. acknowledges support from JSPS KAKENHI grants (JP21H04719 and JP23H04925) and a JST CREST grant (JPMJCR21E6). E.L.A. acknowledges support from the ENCODE Consortium (UM1HG009375) and NHLBI R01HL159176. J.M.E. acknowledges support from NHLBI R01HL159176; the NHGRI IGVF Consortium (UM1HG011972); the Novo Nordisk Foundation (NNF21SA0072102); the NHGRI Genomic Innovator Award (R35HG011324); and the Gordon and Betty Moore and the BASE Research Initiative at the Lucile Packard Children’s Hospital at Stanford University.

## Author Contributions

P.G. and J.M.E. designed and led the study.

P.G., G.M., and B.G.D. conducted CRUDO experiments.

P.G., B.G.D., Y.T., X.S.C., and C.P.F conducted analysis.

J.N., K.M., A.R.G., M.U.S., and A.S.T. generated enhancer-gene pair predictions.

D.T.B. engineered the CRISPRi expressing cell line.

J.R., E.J., C.J.M., A.R.G, L.M.S., and J.M.E. contributed to design and analysis of TAP-seq experiments.

S.S.P.R. and E.L.A. contributed to data interpretation.

S.G.P., N.M., S.S.P.R., R.K., R.M., M.T.K., and E.L.A. facilitated data integration with ENCODE degron experiments.

N.C.D., M.S.S., D.W., S.S.P.R., and E.L.A. facilitated data integration with ENCODE Hi-C experiments.

P.G. and J.M.E. wrote the manuscript with input from all authors.

J.M.E., A.M., and E.S.L. supervised the work.

## Methods

### Enhancer predictions

We used the ABC model to predict enhancer-gene pairs in the presence and absence of cohesin (**Supplementary Table 1**) from measurements of chromatin accessibility (DNase-seq), histone modifications (H3K27ac ChIP–seq), and chromatin conformation (Hi-C) in untreated and 6h auxin treated HCT-116 RAD21-mAID-mClover as previously described^18^. In brief, we call peaks from the chromatin accessibility data using MACS2 with a p-value cut-off of 0.1. We resize the peaks to 500bp centered around the highest signal of the peak, include peaks around the TSS of each gene, and exclude blacklisted regions. For the predictions, we consider all element-gene pairs where the element is within 5Mb of the gene TSS. Next, we compute element activity, by quantifying the chromatin accessibility and H3K27ac ChIP–seq data (both quantile-normalized to the observed distribution) reads within each candidate element data and determine the geometric mean of these two assays. Then, we extract element-promoter contact frequency from chromatin conformation data. Finally, for each element–gene pair, we calculated an ABC score that is the product of activity and contact, normalized by the product of activity and contact for all other elements within 5 Mb of that gene. For the analyses, we included all enhancer-gene pairs meeting the following criteria: the gene has a TPM >=1 in the HCT-116 baseline condition, as measured by PRO-seq^24^, and is non-ubiquitously expressed^18^, with an ABC score of >=0.025693 (this threshold was selected to achieve a 70% recall rate based on previous K562 data^18^). ENCODE-rE2G enhancers were generated and thresholded on the same data set (**Supplementary Table 1)**. Code for analyzing and visualizing HCT116 enhancer predictions in the presence and absence of cohesin is available upon request and will be uploaded to a public repository prior to publication.

### Simulation of cohesin depletion and enhancer perturbation treatment

To simulate the impact of an enhancer perturbation on gene expression in the absence or presence of cohesin we employed a computational model that considers various factors influencing gene regulation. The key parameters of the model include: i) Effect of cohesin removal on overall gene expression, ii) enhancer effect size, iii) effect of cohesin on enhancer function, iv) mRNA half-life, v) CRISPRi induction time point, vi) CRISPRi induction rate, vii) cohesin degradation induction time point, viii) cohesin degradation induction rate, and ix) the measurement timepoint. To assess whether we can distinguish enhancers that rely on cohesin for enhancer function from those that do not, we conducted simulations for distinct cohesin effects on enhancer function with varying values for CRISPRi induction time point, cohesin degradation induction time point, and rate while keeping other parameters constant. Based on these simulations, we determined that 24h of CRISPRi treatment and degradation of cohesin for the last 6h is sufficient to distinguish differential enhancer effects between cohesin present and absent (**Extended Data Fig. 4**). Code for simulations is available upon request and will be uploaded to a public repository prior to publication.

### Target selection

To address how the quantitative contact frequency of an enhancer and its target promoter influence target gene expression levels, we examined pre-existing data on gene expression changes in the absence of cohesin^24^. We analyzed bulk RNA-seq and PRO-seq obtained from untreated and 6-hour auxin treated HCT-116 RAD21-mAID-mClover. We identified ∼150 candidate genes with at least -0.25 and -0.5 log_2_ fold-change in RNA-Seq and PRO-Seq, respectively, and a basal expression of 20 transcripts per million (TPM). From these genes we chose 5 candidate genes with diverse regulatory landscapes based on ABC predicted enhancers spanning a range of linear genomic distances.

### gRNA library design

We designed our gRNA libraries to target the transcription start sites (TSSs) of our genes of interest, candidate enhancers, and non-targeting negative controls. (gRNA sequences are deposited in the IGVF Data Portal under the accession numbers in **Supplementary Table 3a**). Candidate enhancers are 500bp regions centered around the DNase-seq peak of ABC-predicted enhancers (CCND1 pilot CRUDO-FF screen) or within 1Mb of that gene (CRUDO-FF screens, 5Mb for MYC). TSSs are 500bp centered around the DNase peak defined by PRO-seq as the TSS. We designed our gRNAs to be tiled across each element by making a list of every possible target site with an NGG PAM sequence. We then used a previously described algorithm^65^ to rank the target sites by calculating a specificity score based on potential off-target effects. We removed target sites with a score <20 and included up to 30 guides per element for our final pilot gRNA library. We included 400 non-targeting negative controls^66^. For cloning purposes every target sequence had a ‘G’ base added to the 5’ end of the sequence, unless already present.

### CRUDO vector generation

We adapted the CROP-opti vector (Addgene, #106280) and the sgOpti vector (Addgene, #85681) to express a Blasticidin resistance cassette in place of the original Puromycin resistance cassette (CROP-opti-Blast and sgOpti-Blast). To remove the Puromycin resistance cassette we digested the vector with BsiWI (New England Biolabs) and MluI (New England Biolabs) and subsequently purified the linearized vector with 0.5X AMPure XP beads (Beckman Coulter, #A63881). We PCR-amplified the Blasticidin resistance gene from the lenti-dCas-VP64_Blast vector (Addgene, #61425) adding homology arms for Gibson Assembly. We purified the PCR-product with 1X AMPure XP beads and used Gibson Master mix (New England Biolabs) assemble the previously prepared vector backbone and purified Blasticidin PCR-amplified insert following Gibson assembly recommendations. We chemically transformed the assembled product into competent cells (New England Biolabs, #C2987H) and grew the bacteria on LB plates at 30°C overnight. For TAP-seq, we cloned a vector (based on sgOpti-Blast) that would allow us to capture guide transcripts directly with 10x beads. We added the cs1 direct capture sequence^67^ into the gRNA scaffold by digesting out the standard scaffold in sgOpti-Blast with NsiI-HF and EcoRI-HF and then used Gibson assembly to add back in the modified scaffold We selected single colonies to inoculate in liquid culture overnight, prepared the DNA (Machery Nagel, NucleoBond Xtra Midi EF) and screened them by enzyme digestion and Sanger sequencing (plasmid map for CROP-opti-Blast: https://data.igvf.org/documents/4b4305ce-5844-4b84-a3c7-263a1040b050/, plasmid map for sgOpti-Blast-cs1: https://data.igvf.org/documents/6f05cff9-a97a-45f7-9503-ebd5c44873b0/).

### gRNA library cloning

We ordered a PCR-tagged custom oligo pool (Agilent) corresponding to the gRNA library design. We PCR-amplified the gRNA pool to add homology arms for Gibson Assembly. We linearized the CROP-opti-Blast vector by digesting with BsmBI (New England Biolabs) and purified the gRNA pool and vector backbone with 1.5X and 0.7X AMPure XP beads, respectively.

For the Gibson assembly we used 500ng of the purified vector backbone and 70ng of the purified gRNA library with 15μl of 2X Gibson Mix and subsequently purified the assembled product with 0.7X AMPure XP beads eluting in 15μl H2O. We used electroporation to transform the full 15μl of assembled product into 25μl of electrocompetent bacteria (Endura, #60242). We plated an aliquot of the transformed cells on an LB plate in a serial dilution to estimate the number of transformed colonies. We expanded the remaining transformed cells in liquid culture to grow at 30°C overnight. We purified the overnight cultures using an Endotoxin-Free Plasmid Midi Kit (Machery Nagel, NucleoBond Xtra Midi EF). We sequenced gRNAs to verify the library complexity.

### Cell culture

We cultivated HEK293T cells (Takara, #632180) in DMEM media (Thermo Fisher Scientific, #11995065) with 10% HIFBS (Thermo Fisher Scientific, #10082147) added. We maintained a confluency of 20-80% by splitting the cells every two days. We maintained HCT-116 cells in McCoy’s 5A media (Thermo Fisher Scientific, #16600082) supplemented with 2mM L-glutamine (Thermo Fisher Scientific, 25030081), 100U/ml Penicillin with 100μg/ml Streptomycin (Thermo Fisher Scientific, #15140122), and 10% HIFBS at 20-80% confluency by splitting every two days. All cells were stored in an incubator at 37°C and 5% CO2.

### Lentivirus production & transduction

We plated 2.5 million HEK293T cells on a 10cm dish (VWR, #10062880) and 24h later we transfected the cells with 7.2μg of our transfer plasmid as well as 5.4μg of psPAX2 (Addgene, #12260) and 2.16μg of pMD2.G (Addgene, #12259), two lentiviral co-packaging plasmids. We transfected our HEK293T cells using 1150μl Opti-MEM (Thermo Fisher Scientific, #31985070) and 35μl XtremeGene9 (MilliporeSigma, #6365787001) in 10ml of DMEM with 10% HIFBS media. We combined the transfection reagents and incubated these at room temperature for 15 minutes, prior to adding the solution dropwise to the cells. After 16 hours we exchanged the media for fresh DMEM with 10% HIFBS media. At 24 hours post-transfection we harvested viral supernatant for the first time and added fresh DMEM with 10% HIFBS media to the cells. We stored the viral supernatant at 4°C and harvested viral supernatant again at 48 hours post transfection. We combined the harvested virus supernatant for each construct, filtered the supernatant using a vacuum-driven filtration system (MilliporeSigma, #SE1M003M00) and finally concentrated the virus 2X with centrifugal filters (MilliporeSigma #UFC910024). We stored the filtered and concentrated viral supernatant at 4°C unless the time before use extended two weeks, then we divided the virus into 1ml aliquots and snap froze at -80°C.

We transduced HCT-116 cells in 24-well plates with a cell density of 250,000 cells per well and media supplemented with 10μg/ml polybrene (MilliporeSigma #TR1003G). After adding the virus, we infected the cells by centrifuging at 1,200g and 32°C for 45 minutes. 24 hours post-transduction we removed the media containing the virus from the cells and exchanged it for fresh media supplemented with 7.5μg/ml Blasticidin S HCI (Thermo Fisher Scientific, #A1113903) to start the 5-day selection.

### CRISPRi line generation

We generated a CRISPRi inducible HCT-116 RAD21-mAID-mClover cell line by transducing these cells with a plasmid expressing rtTA coupled to a neomycin resistance cassette (Takara, #631363). The cells were then put under selection by supplementing the media with 200μg/ml G418 Sulfate (Thermo Fisher Scientific, #10131035). Positive cells were subsequently transduced with a construct expressing KRAB-dCas9 linked to BFP under a TRE3G promoter (Addgene, #85449). The cells were selected by fluorescent cell sorting (FACS) for BFP by activating the BFP expression with 1μg/ml doxycycline (Thermo Fisher Scientific, #BP26531). Untransduced HCT-116 RAD21-mAID-mClover cells also treated with 1μg/ml doxycycline were used to control for negative BFP expression during cell sorting (**Extended Data Fig. 3b)**.

### Single-guide qPCR validation of promoter targeting guides

We cloned two non-overlapping single guides against the TSSs of the five genes we profiled by CRISPRi-FlowFISH into the CROP-opti-Blast vector (**Extended Data Fig. 13, Supplementary Table 3**). We then infected the single guides into CRISPRi HCT-116 RAD21-mAID-mClover cells at low MOI and selected with Blasticidin as described above. We plated gRNA-expressing stable cell lines at 500,000 cells/ml in two separate plates, both receiving 1 μg/ml doxycycline and one receiving 500μM auxin after 18 hours. 24 hours after doxycycline addition, we extracted RNA from 20,000 cells per experiment in Buffer RLT (Qiagen) using Dynabeads MyOne Silane beads (Thermo Fisher), treated samples with TURBO DNase (Thermo Fisher), and cleaned again with Dynabeads MyOne Silane beads. We used AffinityScript reverse transcriptase (Agilent Technologies, Lexington, MA) and random nonamer primers to convert RNA to cDNA. We performed qPCR using SYBR Green I Master Mix (Roche) with primers for each of the genes and housekeeping control genes and calculated differences using the ΔΔCT method (**Supplementary Table 5**).

### Pooled CRISPRi screen

We transduced doxycycline inducible CRISPRi HCT-116 RAD21-mAID-mClover cells with the gRNA library at a multiplicity of infection (MOI) of about 0.3 and a coverage of 500 transduced cells for each gRNA. After selection we combined all transduced cells and maintained them in 2.5μg/ml Blasticidin at a 500X coverage of the gRNAs at all times.

We expanded the cells to obtain 50 million cells and plated 12.5 million cells each onto four T-75 flasks (Thermo Fisher Scientific, #156499). We added 1μg/ml doxycycline to two of the flasks. 18 hours later we added 500μM auxin (Indole-3-acetic acid, MilliporeSigma, #I37505GA) to one of the doxycycline treated flasks and to one other flask. After a total of 24 hours, we harvested the cells, by aspirating the media, washing the adherent cells with PBS (Thermo Fisher Scientific, #10010023) and incubating the cells with trypsin (Thermo Fisher Scientific, #25200114) for 2 minutes. We resuspended the cells in fresh media and centrifuged the cells at 400g for 5 minutes.

### CRISPRi-FlowFISH screens

We performed CRISPRi-FlowFISH screens as previously described^18^. In brief, we screened two replicates of 10M cells per gene and stained each sample with the PrimeFlow RNA Assay Kit (Thermo Fisher Scientific, #88–18005) for the respective gene with an Alexa Fluor 647 (AF647, ‘Type 1’, **Supplementary Table 4**) probe set and for *RPL13A* to have a positive control housekeeping gene with Alexa Fluor 488 (AF488, ‘Type 4’, **Extended Data Fig. 14a, Supplementary Table 4**). Using the Astrios EQ Sorter (Beckman Coulter, #B25982) we sorted the cells into six bins based on the fluorescence intensity of target genes, normalizing for staining efficiency (**Extended Data Fig. 14b)**. We then performed reverse-crosslinking and gDNA extraction on the collected cells and sequenced the guide RNA integrations as previously described^18^.

### CRISPRi FlowFISH processing

We analyzed CRISPRi-FlowFISH screens as previously described^18^. Briefly, we counted gRNAs in each bin using Bowtie^68^ to map reads to a custom index, normalized gRNA counts in each bin by library size, then used a maximum-likelihood estimation approach to compute the effect size for each gRNA. We used the limited-memory Broyden-Fletcher-Goldfarb-Shanno algorithm (implemented in the R stats4 package) to estimate the most likely log-normal distribution that would have produced the observed guide counts, and the effect size for each gRNA is the mean of its log-normal fit divided by the average of the means from all negative-control gRNAs. As previously described, we scaled the effect size of each gRNA in a screen linearly to account for non-specific target and label probe binding. For each (FlowFISH gene, auxin treatment) pair, we performed single-guide RT-qPCR (**Extended Data Fig. 13**) to determine the absolute effect of knocking these genes down, and used this value to scale our FlowFISH data. Code for processing CRISPRi FlowFISH data is available at https://github.com/EngreitzLab/crispri-flowfish.

### TAP-seq

We performed a modified version of the TAP-seq protocol^39^. We designed a library of 347 gRNAs (gRNA sequences are deposited in the IGVF Data Portal under the accession numbers in **Supplementary Table 3a**) targeting a subset of the screen elements and cloned them as a pool into sgOpti-Blast-cs1 as described above. We designed TAP-seq primers against 74 genes using the pipeline provided^39^, targeting all expressed genes within 1Mb of any of the genes tested by CRISPRi-FlowFISH, as well as control genes. We prepared lentivirus with the pooled gRNA library, infected it at low MOI into CRISPRi HCT-116 RAD21-mAID-mClover cells, and treated with doxycycline and auxin as described above. We loaded 16,000 cells per lane in two technical replicate lanes for no auxin and plus auxin conditions and performed 10x encapsulation and barcoding following manufacturer’s instructions (10x Chromium Next GEM Single Cell 3’ GEM Kit v3.1). We then amplified our target gene panel and gRNA libraries using a mix of primers. We amplified the cDNA through sequential PCRs using a mix of TAP-seq primers for dialing out target genes, and with primers from the 10x Genomics feature barcoding kit to dial out the gRNAs. We confirmed library size using an Agilent TapeStation and sequenced on a Next-seq targeting 20,000 reads per cell for the mRNA and 2,000 reads per cell for the gRNA libraries. Code for processing TAP-seq data is available at https://github.com/argschwind/CRISPRiScreen.

### Element effect size analysis

From Flow-FISH or TAP-seq data we averaged effect sizes of each gRNA across replicates and computed the effect size of an element as the average of all gRNAs targeting that element **(Extended Data Fig. 14c,d)**. We assessed significance using a two-sided t-test comparing the mean effect size of all gRNAs in a candidate element to all negative-control guides. We computed the FDR for elements using the Benjamini-Hochberg procedure and used an FDR threshold of 0.05 to call significant regulatory effects. We identified elements as enhancers if they fulfilled the following criteria: i) a significant effect on gene expression based on a Benjamini-Hochberg FDR correction of <0.05, ii) at least a 5% decrease in target gene expression, iii) measurement values for at least 10 gRNAs, iv) not intra-target genic and not a TSS element, and v) an H3K27ac level above the median of all tested elements. We evaluate differential enhancer effects in the presence and absence of cohesin, we performed a two-sided paired t-test for each element using replicate-averaged gRNA effect sizes of each condition. To determine cohesin dependence, we first determined the effect of each enhancer on its target gene and cap these values at zero for instances where they were less than zero (based on the assumption that an enhancer can either reduce gene expression or not have an effect). We calculated the ‘cohesin dependence’ metric on a per-element basis as the average difference in effect size between gRNAs at baseline and in the absence of cohesin, normalized by the average effect size of gRNAs in the presence of cohesin. The 95% confidence interval for the cohesin dependence was also computed on a per-element basis by using ±1.96 times the standard error of the mean, divided by the average effect size without cohesin presence.

We performed a correlation analysis between our measured CRISPRi enhancer effect sizes and their corresponding predicted ABC scores (**Extended Data Fig. 12**). By fitting a curve to these data, we estimated that an enhancer effect size of 15% corresponds to an ABC score of 0.038 (**Extended Data Fig. 12).**

Code for analyzing and visualizing cohesin CRUDO-FlowFISH or -TAP-seq data is available upon request and will be uploaded to a public repository prior to publication.

## Extended Data Figures

**Extended Data Fig. 1.**
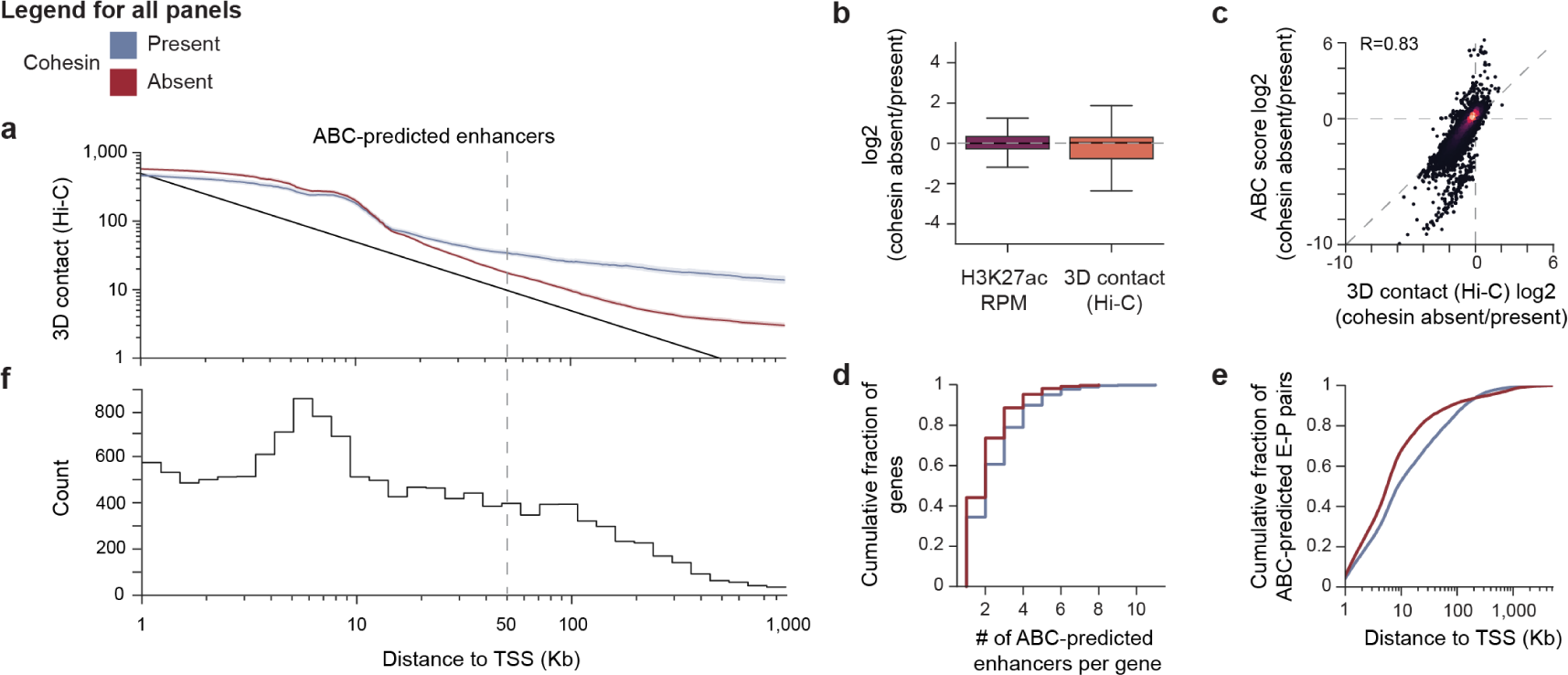
Intrinsic enhancer activity and contact frequency in the ABC model. **a**, Enhancer-promoter 3D contact frequency (Hi-C SCALE-normalized counts, 5-Kb resolution^35^) in the presence (blue, no auxin, Hi-C depth = 1.1 billion contacts) or absence (red, plus 6 hours auxin, Hi-C depth = 1.5 billion contacts) of cohesin, as a function of distance from enhancer to target promoter. Lines: Rolling average for all enhancer-gene regulatory interactions predicted by ABC smoothed over 1,000 data point windows. Black line: Enhancer-promoter contact frequency predicted by the inverse power law model. **b,** Changes in H3K27ac levels and 3D contact frequency (Hi-C SCALE-normalized counts, 5-Kb resolution) between cohesin absent and present for all enhancer-promoter regulatory interactions predicted by ABC (n=14,274). Boxes: median and interquartile range. Whiskers: full distribution, except for points determined “outliers” using the interquartile range. **c,** Change in ABC score between cohesin present and absent in log_2_ as a function of the change in Hi-C contact frequency (Hi-C counts at a 5Kb resolution) between cohesin present and absent in log_2_ for all enhancer-promoter regulatory interactions predicted by ABC (n=14,274). **d,** Considering genes that have at least one predicted ABC enhancer, cumulative distribution plot of the number of ABC enhancers per gene in the presence of cohesin (blue, n=5,850) and absence of cohesin (red, n=5,190). **e,** Considering genes that have at least one predicted ABC enhancer, cumulative distribution plot of the distance from enhancer to target gene TSS for each ABC-predicted enhancer-promoter pair in HCT116 cells in the presence of cohesin (blue, n=14,274) and the absence of cohesin (red, n=10,514). **f**, Considering genes that have at least one predicted ABC enhancer, count of ABC-predicted enhancer-promoter pairs in HCT116 cells in the presence of cohesin (n=14,274, as a function of distance from enhancer to target promoter.

**Extended Data Fig. 2.**
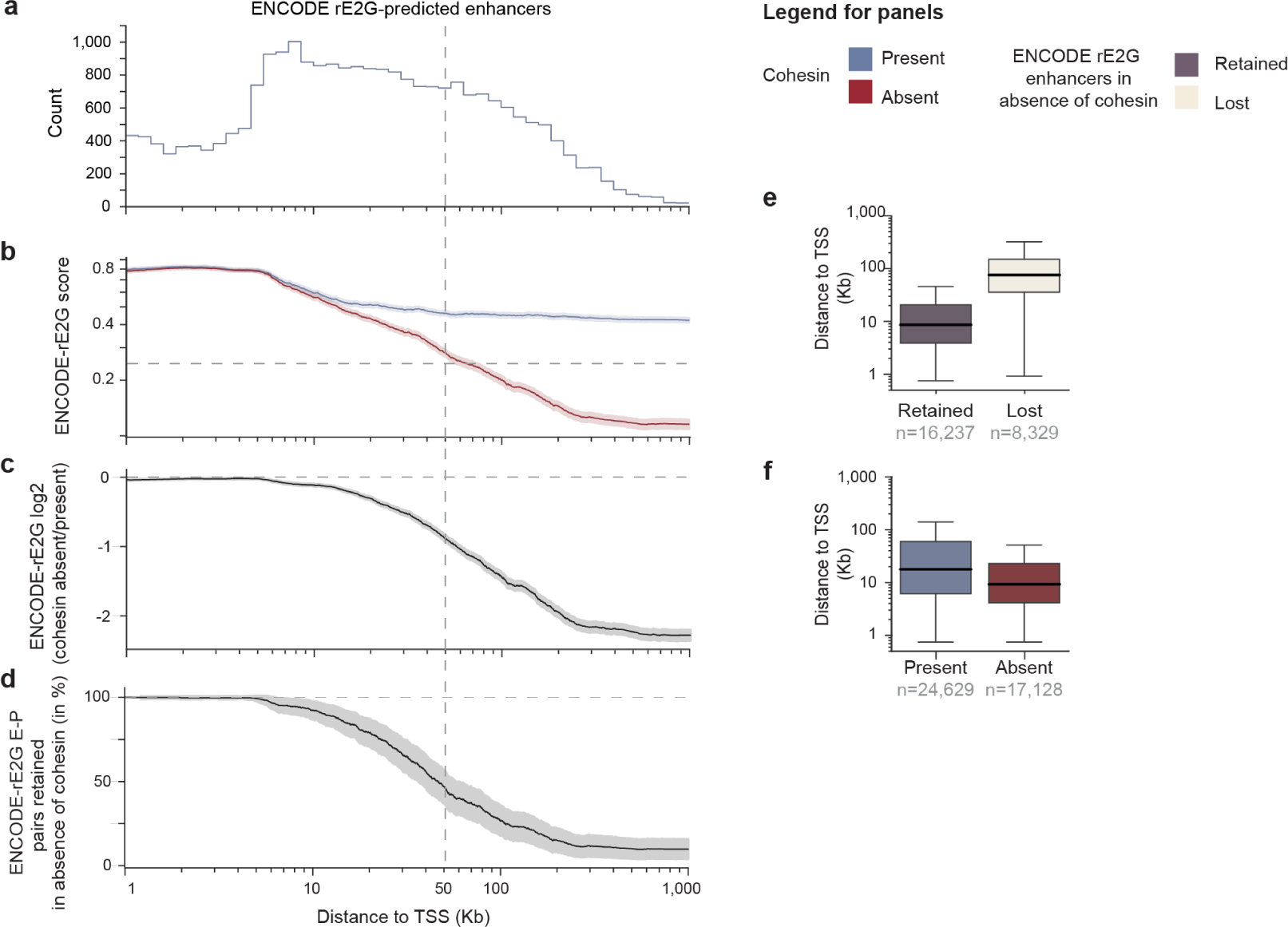
Summary ABC enhancer predictions statistics for the HCT116 cell line in the presence and absence of cohesin. **a**, Considering genes that have at least one predicted ENCODE-rE2G enhancer, count of ENCODE-rE2G-predicted enhancer-promoter pairs in HCT116 cells in the presence of cohesin (n=24,629), as a function of distance from enhancer to target promoter. **b,** Similar to **a**, showing ENCODE-rE2G score in the presence (blue, no auxin) or absence (red, plus 6 hours auxin) of cohesin, as a function of distance from enhancer to target promoter. Lines: Rolling average for all enhancer-gene regulatory interactions predicted by ABC in the cohesin present condition (n=24,629),) smoothed over 1,000 data point windows. Shading: 95% c.i. of the rolling average. **c,** Similar to **a**, showing a log_2_ fold-change in ENCODE-rE2G score between the absence and presence of cohesin as a function of distance. **d,** Similar to **a**, showing percentage of enhancer-promoter regulatory interactions predicted by ENCODE-rE2G in the cohesin present condition that also pass the ENCODE-rE2G score threshold (0.243) in the absence of cohesin (retained enhancers, n=16,237), as a function of distance. **e,** Distribution of distances from enhancer to target promoter for all enhancer-promoter ENCODE-rE2G-predicted regulatory interactions retained (dark gray, n=16,237) or lost (cream, n=8,329) in the absence of cohesin. **f,** Distribution of distances from enhancer to target promoter for all enhancer-promoter regulatory interactions predicted by ENCODE-rE2G in the presence (blue, n=24,629) or absence of cohesin (red, n=17,128).

**Extended Data Fig. 3.**
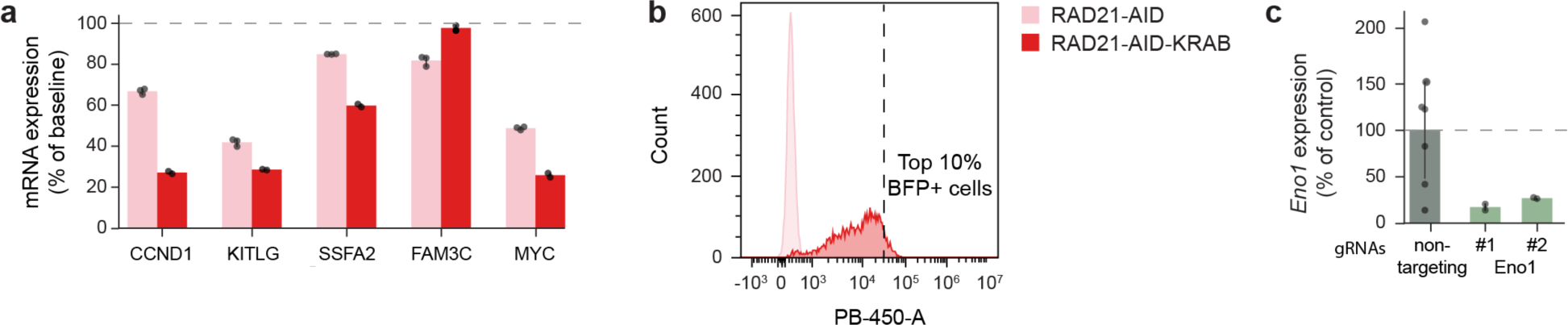
Validating the cohesin depletion cell line additionally expressing the CRISPRi machinery. **a**, mRNA expression levels in the absence of cohesin (6h of auxin treatment) as measured by bulk RNA-seq in the HCT116 cohesin depletion line (rose) and the further engineered cohesin depletion with CRISPRi (red) for the CRUDO-FF screen genes. Error bars represent the 95% c.i.s. **b**, FACS showing dox inducibility of HCT-116 RAD21-mAiD-mClover-KRAB-dCAS9 after sorting but before the screen. HCT-116 RAD21-mAiD-mClover control cells (rose) and HCT-116 RAD21-mAiD-mClover-KRAB-dCAS9 (red) cells were treated with Doxycycline for 24h and the analyzed. Live cells were selected using FSC-A/SSC-A, doublets were similarly removed with FSC-H/FSC-W and SSC-H/SSC-W, and the top 10% of the HCT-116 RAD21-mAiD-mClover-KRAB-dCAS9 BFP-positive peak (PB450-A) were sorted. **c,** qPCR for *ENO1* in the HCT116 cohesin depletion CRISPRi line for negative control guides (non-targeting) and two ENO1-targeting guides (ENO1-1, ENO1-2). Error bars represent the 95% c.i..

**Extended Data Fig. 4.**
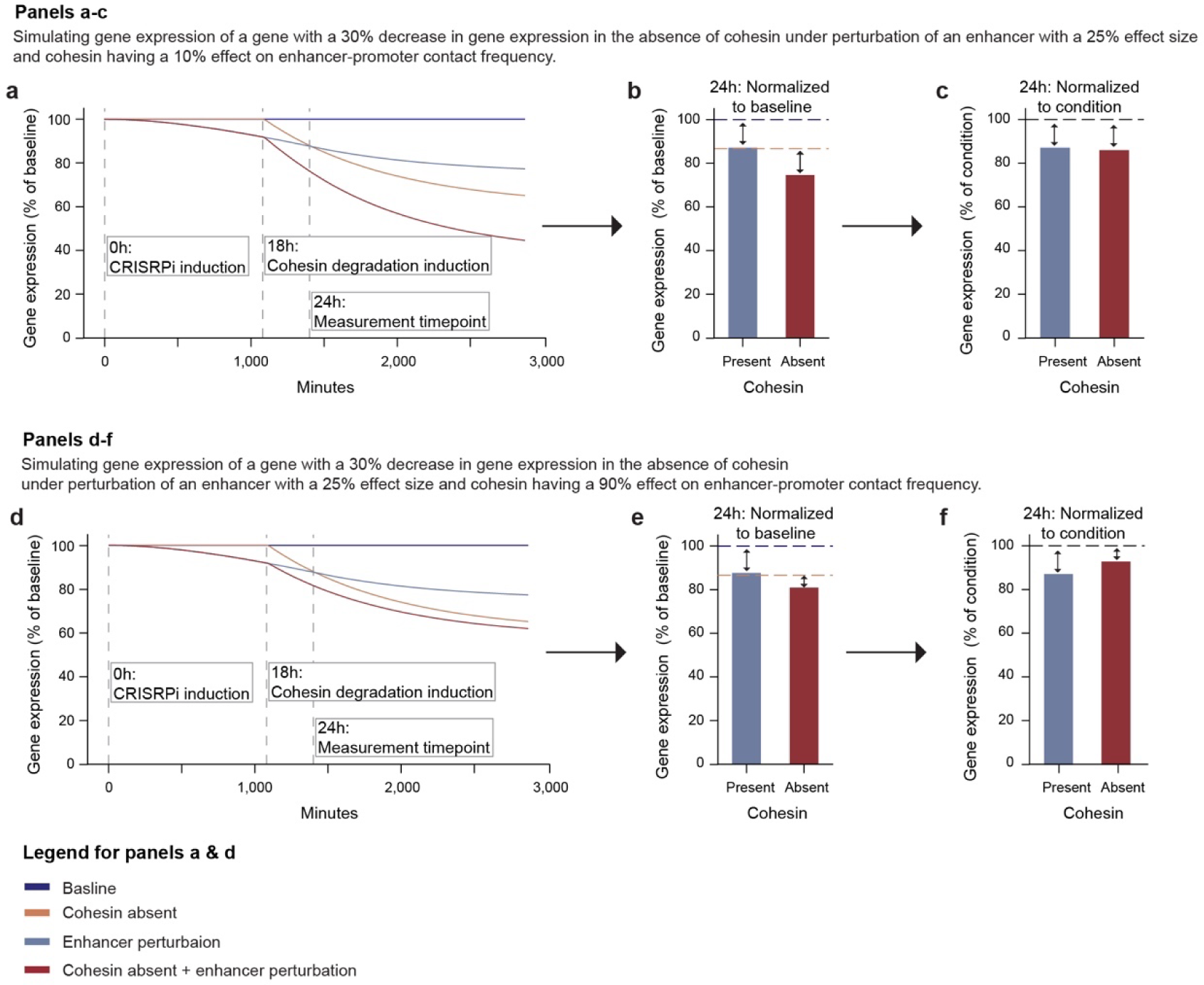
Simulation of cohesin depletion and enhancer perturbation treatment. **a**, 48h gene expression simulation for four conditions - Baseline (dark blue), Cohesin absent (peach), Enhancer perturbation (light blue), and Cohesin absent plus enhancer perturbation (red) - under perturbation of an enhancer induced at 0h with a 25% effect on its target gene that has an mRNA half-life of 6 hours and loses 30% of gene expression in the absence of cohesin after 6h with a 10% effect of cohesin on enhancer-promoter contact frequency. 24h zoom in of the perturbation as percent of the baseline (**b**) and the respective cohesin condition (**c**). **d,** Similar to **a**, but with a 90% effect of cohesin on enhancer-promoter contact. frequency. **e**, Similar to **b**, with a 90% effect of cohesin on enhancer-promoter contact frequency. **f,** Similar to **c**, with a 90% effect of cohesin on enhancer-promoter contact frequency.

**Extended Data Fig. 5.**
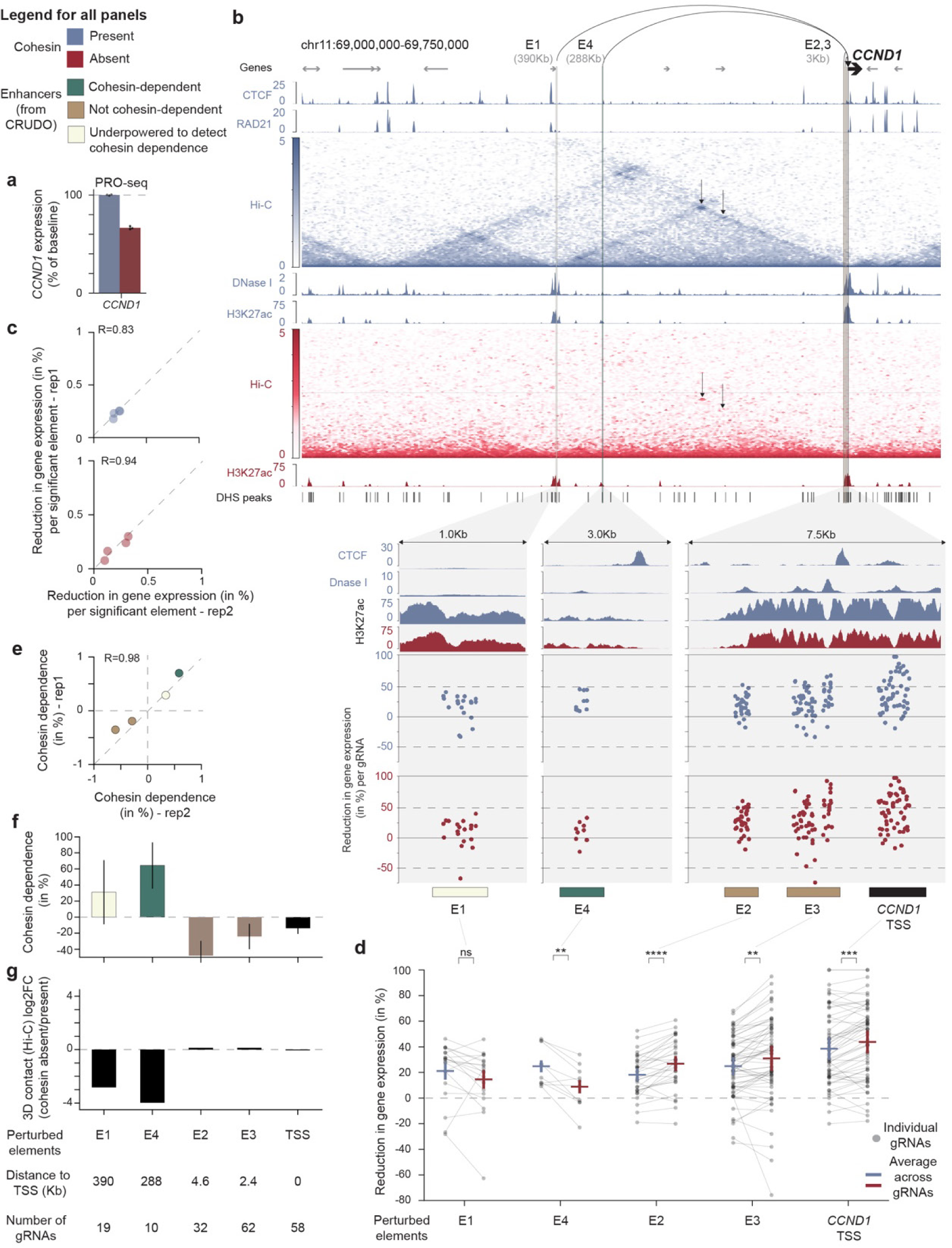
Browser snapshot of the CCND1 locus with CRUDO-FF screen effects. **a**, *CCND1* transcript levels in the presence (blue, 6h of DMSO treatment) or absence (red, 6h of auxin treatment) of cohesin, as measured by PRO-seq. Error bars: 95% c.i. across 3 biological replicates. **b,** *CCND1* locus showing DNase I hypersensitivity, CTCF ChIP-seq, RAD21 ChIP-seq, SCALE-normalized Hi-C at 5Kb resolution (Hi-C depth = 1.1 (no auxin) and 1.5 (plus 6 hours auxin) billion contacts)^35^, and H3K27ac ChIP-seq in the presence (blue) and/or absence (red) of cohesin. Bars at bottom show locations of all elements tested in the CRUDO-FF experiment. Zoom-ins show all elements that had a significant effect on *CCND1* expression in cohesin present condition. Bars at bottom show genomic locations of elements identified in the CRUDO-FF screen (teal: cohesin-dependent enhancers; taupe: not cohesin-dependent enhancers; ivory: underpowered to detect cohesin dependence enhancers; black: TSS). Dots show the measured effects on *CCND1* expression for individual gRNAs targeting each element (% reduction in gene expression, normalized to the mean of all non-targeting gRNAs within each condition). **c**, Scatterplot shows correlation of effects on gene expression between two biological replicate experiments in the cohesin present (blue, top panel) and cohesin absent (red, bottom panel) conditions. Dots: all 4 elements that had significant effects on gene expression in the cohesin present condition. **d,** CRUDO-FF measured effect sizes on *CCND1* expression for individual gRNAs in the presence (blue) or absence of cohesin (red). Horizontal lines and error bars: mean (+/- 95% c.i.) of effect sizes for each element. ns: *P_BenjaminiHochberg_* > 0.05, **: *P_BenjaminiHochberg_* < 0.01, ***: *P_BenjaminiHochberg_* < 0.001, ****: *P_BenjaminiHochberg_* < 0.0001 (gRNA-based paired 2-sample t-test). **e,** Similar to panel **c,** showing correlation of cohesin dependence between two biological replicate experiments. **f**, The level of cohesin dependence for each element, describing the change in the effect of an element on gene expression between the cohesin present versus absent conditions. Bar: mean across gRNAs for each element. Error bars: 95% c.i.. **g**, Log_2_ fold-change in 3D contact frequencies (Hi-C SCALE-normalized counts, 5Kb resolution, Hi-C depth = 1.1 (no auxin) and 1.5 (plus 6 hours auxin) billion contacts^35^) between the absence and presence of cohesin.

**Extended Data Fig. 6.**
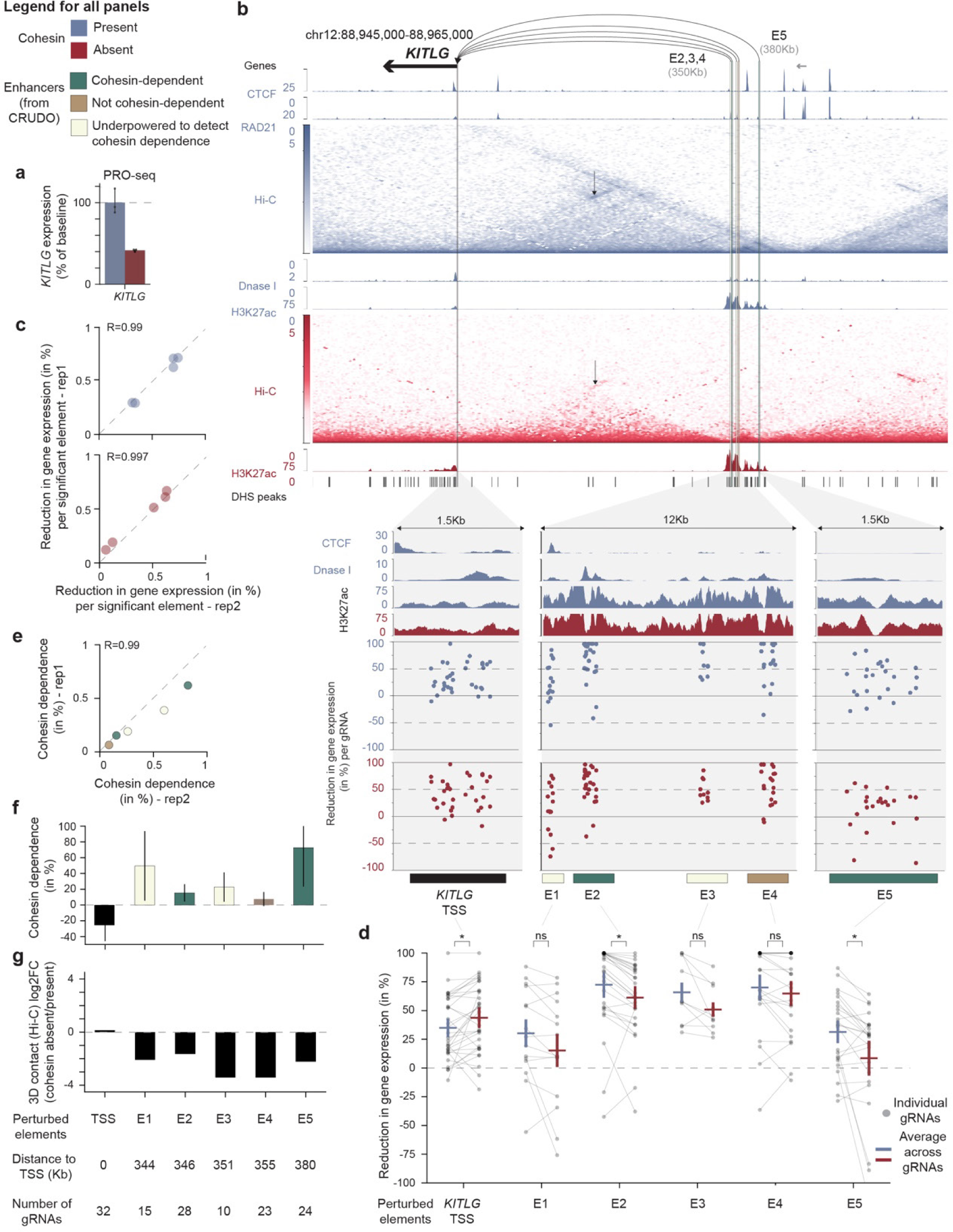
Browser snapshot of the KITLG locus with CRUDO-FF screen effects. **a**, *KITLG* transcript levels in the presence (blue, 6h of DMSO treatment) or absence (red, 6h of auxin treatment) of cohesin, as measured by PRO-seq. Error bars: 95% c.i. across 3 biological replicates. **b,** *KITLG* locus showing DNase I hypersensitivity, CTCF ChIP-seq, RAD21 ChIP-seq, SCALE-normalized Hi-C at 5Kb resolution (Hi-C depth = 1.1 (no auxin) and 1.5 (plus 6 hours auxin) billion contacts)^35^, and H3K27ac ChIP-seq in the presence (blue) and/or absence (red) of cohesin. Bars at bottom show locations of all elements tested in the CRUDO-FF experiment. Zoom-ins show all elements that had a significant effect on *KITLG* expression in cohesin present condition. Bars at bottom show genomic locations of elements identified in the CRUDO-FF screen (teal: cohesin-dependent enhancers; taupe: not cohesin-dependent enhancers; ivory: underpowered to detect cohesin dependence enhancers; black: TSS). Dots show the measured effects on *KITLG* expression for individual gRNAs targeting each element (% reduction in gene expression, normalized to the mean of all non-targeting gRNAs within each condition). **c**, Scatterplot shows correlation of effects on gene expression between two biological replicate experiments in the cohesin present (blue, top panel) and cohesin absent (red, bottom panel) conditions. Dots: all 5 elements that had significant effects on gene expression in the cohesin present condition. **d,** CRUDO-FF measured effect sizes on *KITLG* expression for individual gRNAs in the presence (blue) or absence of cohesin (red). Horizontal lines and error bars: mean (+/- 95% c.i.) of effect sizes for each element. ns: *P_BenjaminiHochberg_* > 0.05, \**P_BenjaminiHochberg_* < 0.05,(gRNA-based paired 2-sample t-test). **e,** Similar to panel **c,** showing correlation of cohesin dependence between two biological replicate experiments. **f**, The level of cohesin dependence for each element, describing the change in the effect of an element on gene expression between the cohesin present versus absent conditions. Bar: mean across gRNAs for each element. Error bars: 95% c.i.. **g**, Log_2_ fold-change in 3D contact frequencies (Hi-C SCALE-normalized counts, 5Kb resolution, Hi-C depth = 1.1 (no auxin) and 1.5 (plus 6 hours auxin) billion contacts^35^) between the absence and presence of cohesin.

**Extended Data Fig. 7.**
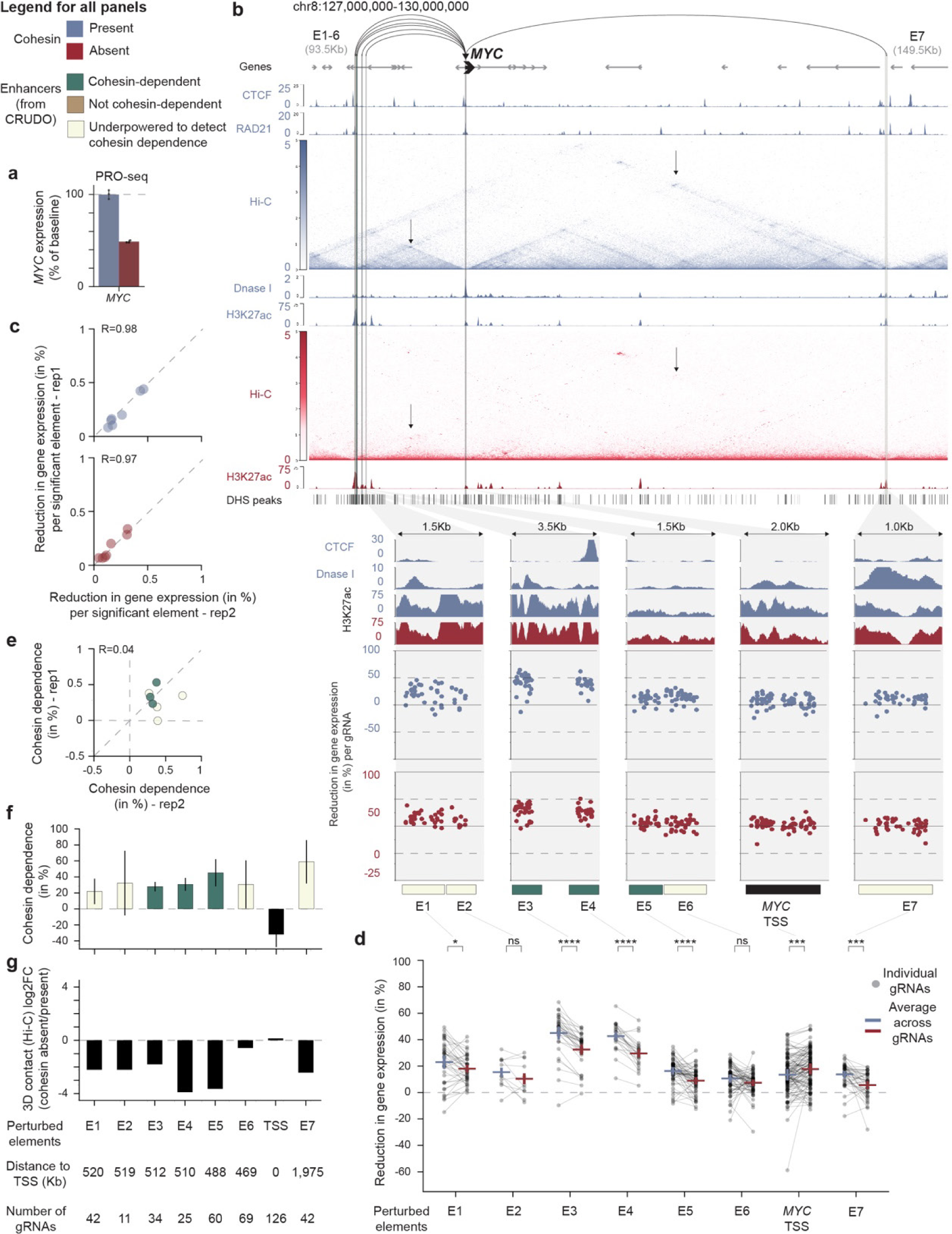
Browser snapshot of the MYC locus with CRUDO-FF screen effects. **a**, *MYC* transcript levels in the presence (blue, 6h of DMSO treatment) or absence (red, 6h of auxin treatment) of cohesin, as measured by PRO-seq. Error bars: 95% c.i. across 3 biological replicates. **b,** *MYC* locus showing DNase I hypersensitivity, CTCF ChIP-seq, RAD21 ChIP-seq, SCALE-normalized Hi-C at 5Kb resolution (Hi-C depth = 1.1 (no auxin) and 1.5 (plus 6 hours auxin) billion contacts)^35^, and H3K27ac ChIP-seq in the presence (blue) and/or absence (red) of cohesin. Bars at bottom show locations of all elements tested in the CRUDO-FF experiment. Zoom-ins show all elements that had a significant effect on *MYC* expression in cohesin present condition. Bars at bottom show genomic locations of elements identified in the CRUDO-FF screen (teal: cohesin-dependent enhancers; taupe: not cohesin-dependent enhancers; ivory: underpowered to detect cohesin dependence enhancers; black: TSS). Dots show the measured effects on *MYC* expression for individual gRNAs targeting each element (% reduction in gene expression, normalized to the mean of all non-targeting gRNAs within each condition). **c**, Scatterplot shows correlation of effects on gene expression between two biological replicate experiments in the cohesin present (blue, top panel) and cohesin absent (red, bottom panel) conditions. Dots: all 7 elements that had significant effects on gene expression in the cohesin present condition. **d,** CRUDO-FF measured effect sizes on *MYC* expression for individual gRNAs in the presence (blue) or absence of cohesin (red). Horizontal lines and error bars: mean (+/- 95% c.i.) of effect sizes for each element. ns: *P_BenjaminiHochberg_* > 0.05, *: *P_BenjaminiHochberg_* < 0.05, ***: *P_BenjaminiHochberg_* < 0.001, ****: *P_BenjaminiHochberg_* < 0.0001 (gRNA-based paired 2-sample t-test). **e,** Similar to panel **c,** showing correlation of cohesin dependence between two biological replicate experiments. **f**, The level of cohesin dependence for each element, describing the change in the effect of an element on gene expression between the cohesin present versus absent conditions. Bar: mean across gRNAs for each element. Error bars: 95% c.i.. **g**, Log_2_ fold-change in 3D contact frequencies (Hi-C SCALE-normalized counts, 5Kb resolution, Hi-C depth = 1.1 (no auxin) and 1.5 (plus 6 hours auxin) billion contacts^35^) between the absence and presence of cohesin.

**Extended Data Fig. 8.**
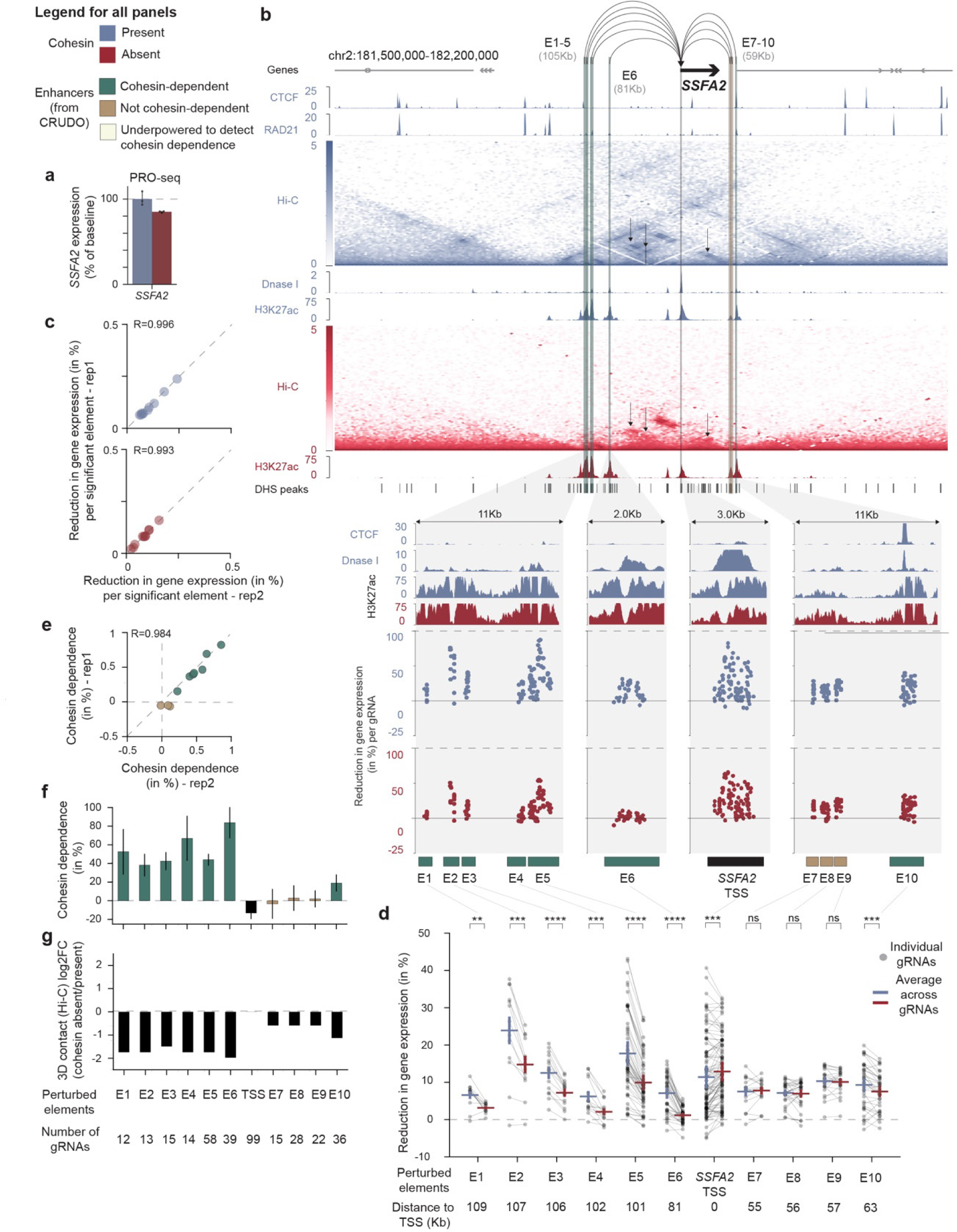
Browser snapshot of the SSFA2 locus with CRUDO-FF screen effects. **a**, *SSFA2* transcript levels in the presence (blue, 6h of DMSO treatment) or absence (red, 6h of auxin treatment) of cohesin, as measured by PRO-seq. Error bars: 95% c.i. across 3 biological replicates. **b,** *SSFA2* locus showing DNase I hypersensitivity, CTCF ChIP-seq, RAD21 ChIP-seq, SCALE-normalized Hi-C at 5Kb resolution (Hi-C depth = 1.1 (no auxin) and 1.5 (plus 6 hours auxin) billion contacts)^35^, and H3K27ac ChIP-seq in the presence (blue) and/or absence (red) of cohesin. Bars at bottom show locations of all elements tested in the CRUDO-FF experiment. Zoom-ins show all elements that had a significant effect on *SSFA2* expression in cohesin present condition. Bars at bottom show genomic locations of elements identified in the CRUDO-FF screen (teal: cohesin-dependent enhancers; taupe: not cohesin-dependent enhancers; ivory: underpowered to detect cohesin dependence enhancers; black: TSS). Dots show the measured effects on *SSFA2* expression for individual gRNAs targeting each element (% reduction in gene expression, normalized to the mean of all non-targeting gRNAs within each condition). **c**, Scatterplot shows correlation of effects on gene expression between two biological replicate experiments in the cohesin present (blue, top panel) and cohesin absent (red, bottom panel) conditions. Dots: all 10 elements that had significant effects on gene expression in the cohesin present condition. **d,** CRUDO-FF measured effect sizes on *SSFA2* expression for individual gRNAs in the presence (blue) or absence of cohesin (red). Horizontal lines and error bars: mean (+/- 95% c.i.) of effect sizes for each element. ns: *P_BenjaminiHochberg_* > 0.05, **: *P_BenjaminiHochberg_* < 0.01, ***: *P_BenjaminiHochberg_* < 0.001, ****: *P_BenjaminiHochberg_* < 0.0001 (gRNA-based paired 2-sample t-test). **e,** Similar to panel **c,** showing correlation of cohesin dependence between two biological replicate experiments. **f**, The level of cohesin dependence for each element, describing the change in the effect of an element on gene expression between the cohesin present versus absent conditions. Bar: mean across gRNAs for each element. Error bars: 95% c.i.. **g**, Log_2_ fold-change in 3D contact frequencies (Hi-C SCALE-normalized counts, 5Kb resolution, Hi-C depth = 1.1 (no auxin) and 1.5 (plus 6 hours auxin) billion contacts^35^) between the absence and presence of cohesin.

**Extended Data Fig. 9.**
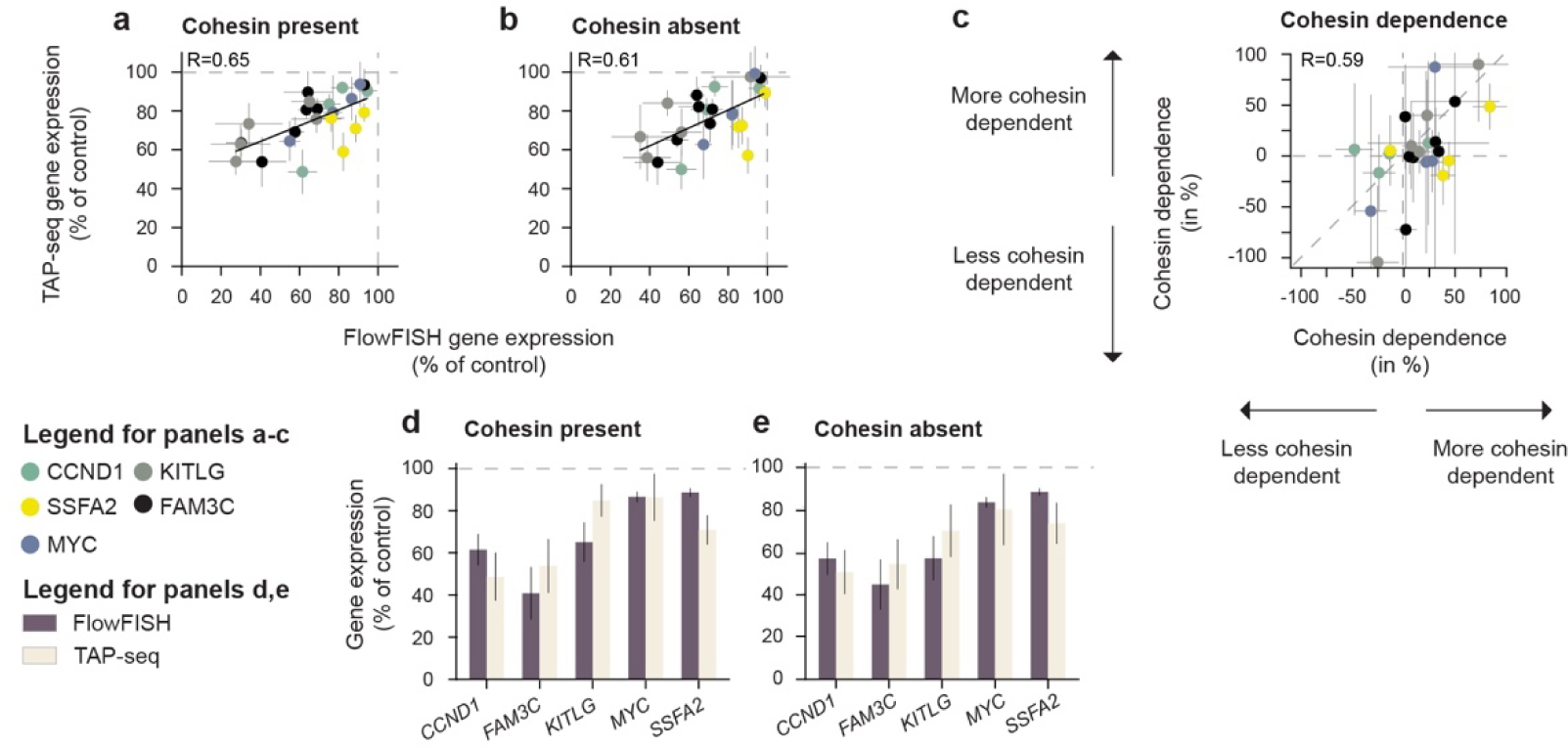
Gene expression measurement correlation between FlowFISH and TAP-seq. **a**, Scatterplot shows correlation of effect size estimates of perturbing a subset of regulatory elements tested in the CRUDO-FF screen in the presence of cohesin as measured by FlowFISH and TAP-seq. Colors represent target genes, Sage: CCND1, Gray: KITLG, Yellow: SSFA2, Black: FAM3C. Blue: Myc. **b,** Similar to **a**, but in the absence of cohesin. **c,** Similar to **a**, but of cohesin dependence as determined by FlowFISH and TAP-seq. **d,** Effect size estimates of perturbing the TSS of the 5 CRUDO-FF screen genes in the presence of cohesin as measured by FlowFISH (dark grey) and TAP-seq (cream). **e,** Similar to **d**, but in the absence of cohesin.

**Extended Data Fig. 10.**
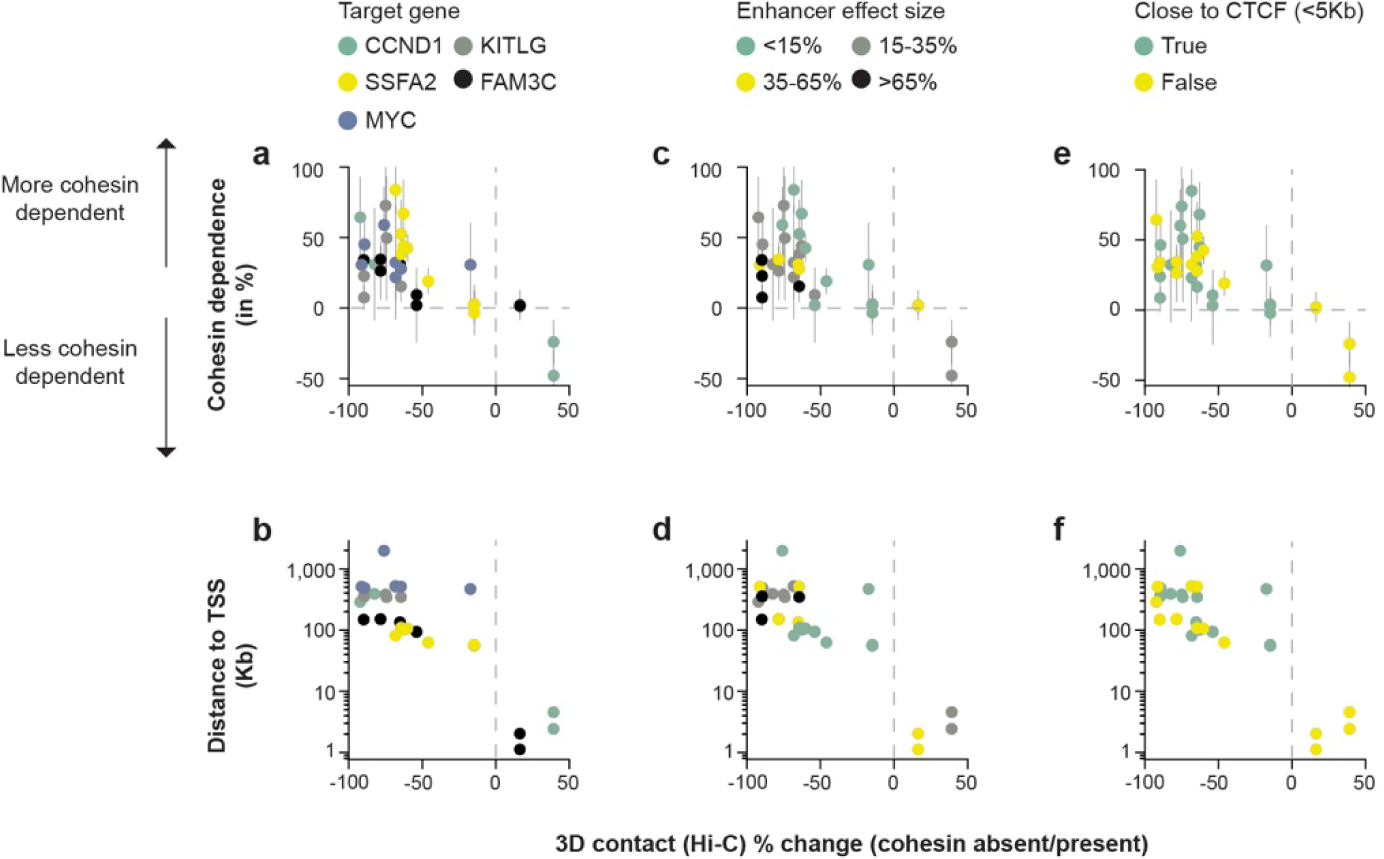
CRUDO-FF screen results colored by features. **a**, Comparison of cohesin dependence with percent change in 3D contacts (Hi-C SCALE-normalized counts, 5-Kb resolution, Hi-C depth = 1.1 (no auxin) and 1.5 (plus 6 hours auxin) billion contacts^35^) between the cohesin absent versus present conditions across all enhancers (n=34). Colors represent target genes, Sage: CCND1, Gray: KITLG, Yellow: SSFA2, Black: FAM3C. Blue: Myc. **b,** Genomic distance (log_10_ scale) from enhancer to target gene TSS versus the fold-change in 3D contacts (Hi-C SCALE-normalized counts, 5-Kb resolution, Hi-C depth = 1.1 (no auxin) and 1.5 (plus 6 hours auxin) billion contacts^35^) between the absence and presence of cohesin. Colors represent target genes, Sage: CCND1, Gray: KITLG, Yellow: SSFA2, Black: FAM3C. Blue: Myc. **c,** Similar to **a**, but colors represent baseline enhancer effect sizes, Sage: <15%, Gray: 15-35%, Yellow: 35-65%, Black: >65%. **d,** Similar to **b**, but colors represent baseline enhancer effect sizes, Sage: <15%, Gray: 15-35%, Yellow: 35-65%, Black: >65%. **e,** Similar to **a**, but colors represent proximity to a CTCF binding site, Sage: <5Kb, Yellow: >5Kb. **f,** Similar to **b**, but colors represent proximity to a CTCF binding site, Sage: <5Kb, Yellow: >5Kb.

**Extended Data Fig. 11.**
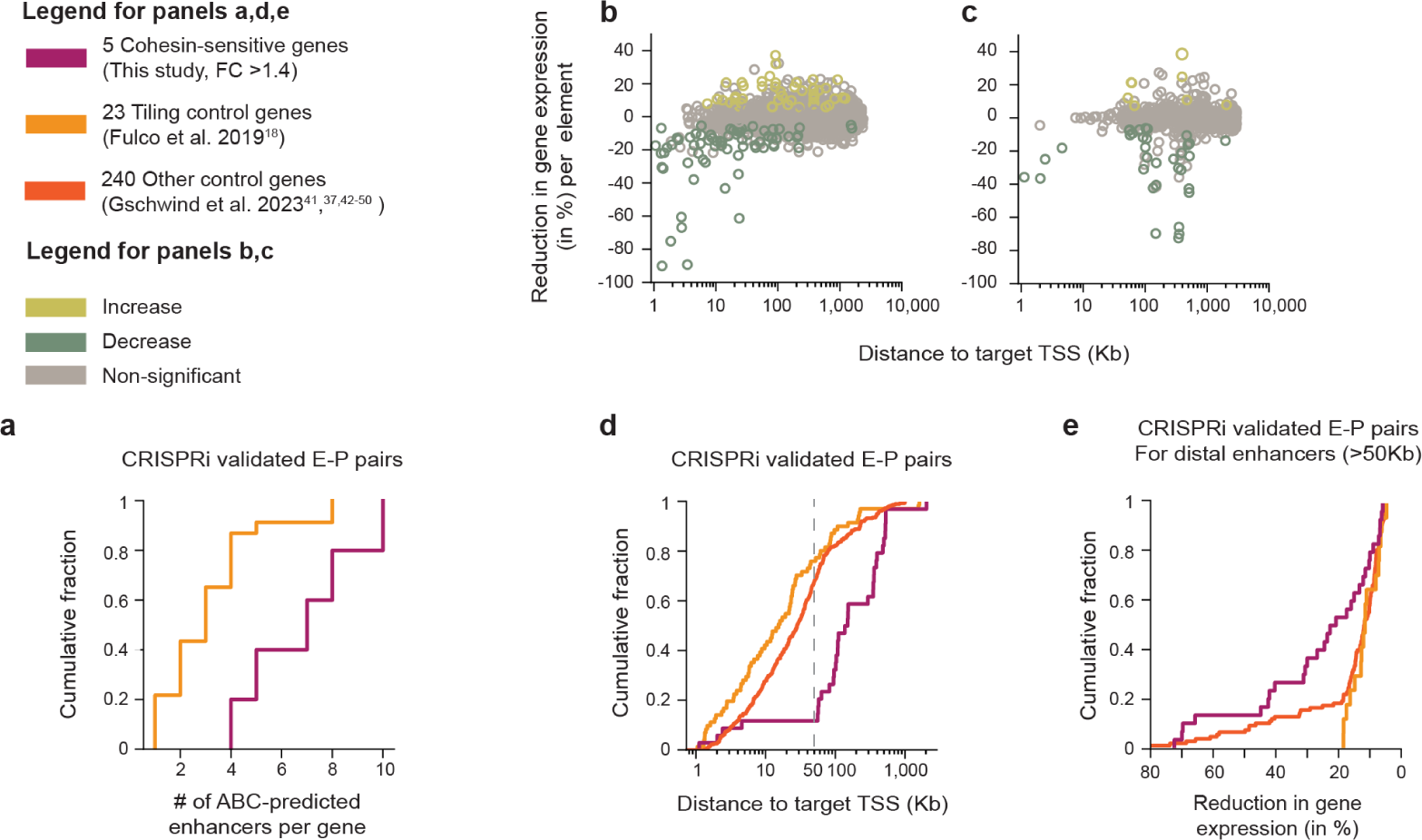
Enhancer property comparison between cohesin-dependent and other genes. **a**, Cumulative distribution plot of the number of enhancers per gene from comprehensive CRISPRi tiling data for the 5 cohesin-dependent genes in HCT116 cells (purple, n=34 this study) and for 23 “Tiling control” genes we previously studied in K562 cells, which were selected without respect to their cohesin dependence (peach, n=71;^18^). **b,** Enhancer effect sizes (as estimated by its _perturbation_) as a function of distance from element to target promoter, for all CRUDO-FF screen tested element-promoter pairs (n=1,086)**. c,** Enhancer effect sizes (as estimated by its perturbation) as a function of distance from element to target promoter, for all tested element-promoter pairs in the K562 “Tiling control” data set (n=4,914,^18^). **d,** Cumulative distribution plot of the genomic distance from enhancer to target gene TSS for each significant pair from comprehensive CRISPRi tiling data for the 5 cohesin-dependent genes in HCT116 cells (purple, n=34 this study), for the 23 “Tiling control” genes^18^, and for the 240 “Other control” genes compiled previously (Gschwind et al.^38^, which were selected without respect to their cohesin dependence and effects were measured using techniques other than CRISPRi-FF (orange, n=240;^39,42–50^). **e,** Cumulative distribution plot of the effect size for each distal enhancer (>50Kb distance to TSS) in the CRISPRi validated enhancer-promoter pairs from our CRUDO-FF dataset (purple, n=30), the “Tiling control” dataset^18^ (peach, n=71), and the “Other control” datset^38,39,42–50^ (orange, n=347).

**Extended Data Fig. 12.**
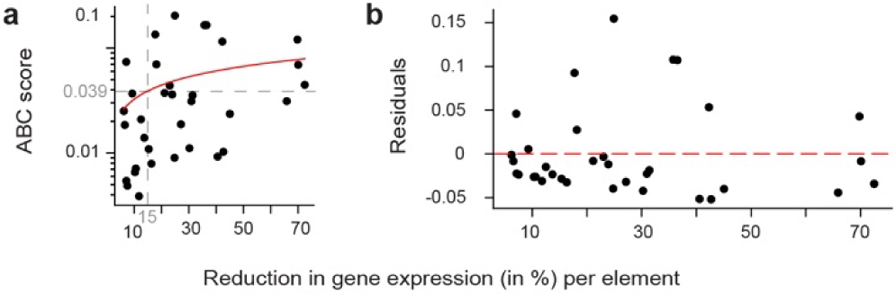
Correlation between CRUDO-FF enhancer effect sizes and ABC score. **a**, ABC score as a function of FlowFISH measured enhancer effect sizes for all CRUDO-FF screen identified enhancer-promoter pairs. The red line represents a power law fit. **b,** ABC score residuals (power law based predicted ABC score subtracted from the actual ABC score) as a function of FlowFISH measured enhancer effect sizes for all CRUDO-FF screen identified enhancer-promoter pairs.

**Extended Data Fig. 13.**
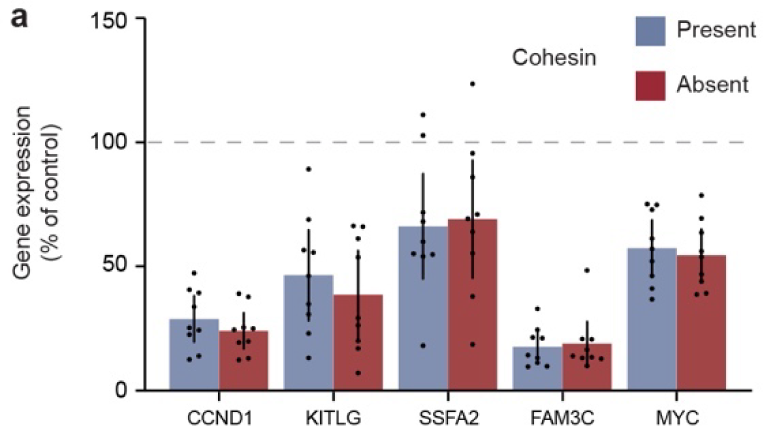
Individual guide qPCR knockdown for CRUDO-FF scaling. **a**, mRNA expression levels in the presence and absence of cohesin (6h of auxin treatment) as measured by qPCR for each of the CRUDO-FF screen genes in cells expressing individual gRNAs targeting the respective TSS for scaling of the CRUDO-FF effect size estimation. Error bars represent the 95% c.i.s.

**Extended Data Fig. 14.**
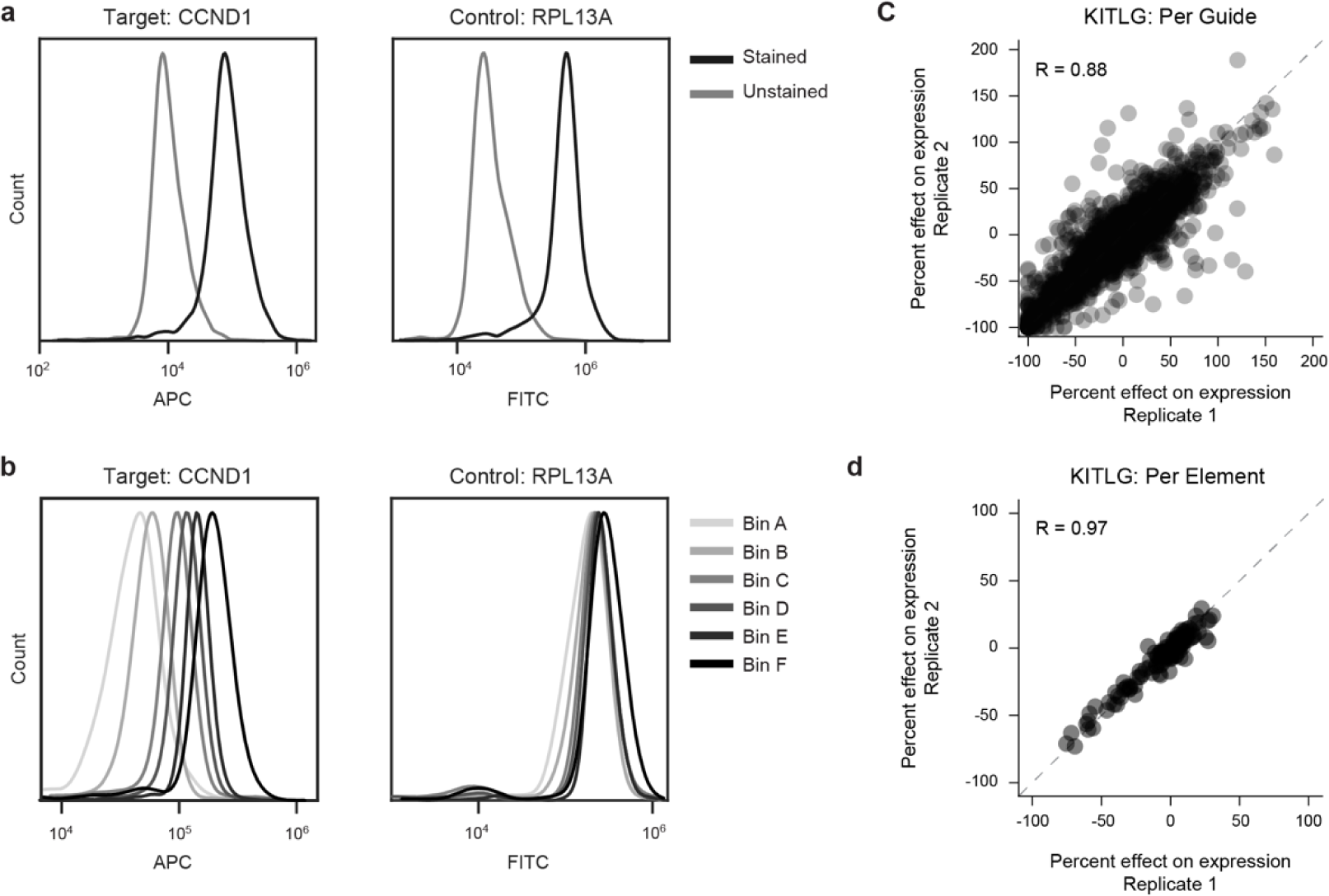
FlowFISH quality controls. **a**, Flow cytometry signal for cells with or without label probe for both target and control genes. Density plots have been normalized to the mode. **b,** Flow cytometry signal for cells after sorting by expression on CCND1 levels (with bin A having the cells with the lowest CCND1 expression and bin F the cells with the highest). Density plots have been normalized to the mode. **c,** Scatterplot of correlation of guide-level effects as estimated from two separate biological replicate experiments. **d,** Scatterplot of correlation of element-level effects as estimated from two separate biological replicate experiments.

